# Selectivity and evolution of Aqp10 in solute permeability influenced by pore molecular weight

**DOI:** 10.1101/2024.03.16.584933

**Authors:** Ayumi Nagashima, Kazutaka Ushio, Hidenori Nishihara, Jin Akimoto, Akira Kato, Tadaomi Furuta

## Abstract

Aqp10 is an aquaglyceroporin that transports not only water but also uncharged low-molecular-weight compounds. We previously demonstrated the evolution of solute permeability in Aqp10 paralogs and showed that the urea and boric acid permeabilities of Aqp10.2 were much weaker than those of Aqp10.1 and plesiomorphic Aqp10s. However, the molecular mechanism responsible for the weak permeability of Aqp10.2 to urea and boric acid remains unclear. Herein, we present a novel hypothesis that explains the solute selectivity of Aqp10. We deduced the ancestral sequences of Aqp10.1 and Aqp10.2 paralogs via molecular phylogenetic analysis. Constructed structural models of these sequences revealed that both the well known amino acid site at position 3 and the sum of molecular weights of the four amino acid sites in the ar/R region were important for the formation of the Aqp10 selectivity filter. Site-directed mutagenesis revealed that a decrease in the sum of the molecular weights of the four amino acid sites enhanced the Aqp10 permeability to urea and boric acid. Based on this, we proposed a model in which the presence of two or more bulky amino acids in the ar/R region, which increases the sum of the molecular weights of amino acids in the ar/R region, was essential for the formation of a filter that limited urea and boric acid transport. Our results outline the molecular mechanism by which Aqp10.2 acquired a selectivity filter during evolution and provide structural insights into the narrowly tuned filter responsible for the solute selectivity of aquaglyceroporins.

**Significance Statement:** In aquaglyceroporins, particularly Aqp10s, the urea and boric acid permeabilities of Aqp10.2 paralogs are much weaker than those of plesiomorphic Aqp10s. Here, we deduced the ancestral sequences of Aqp10.1 and Aqp10.2 via molecular phylogenetic analysis. Structural models of these sequences revealed that the sum of the four amino acid site molecular weights in the ar/R region, if more than one is bulky, contributed to the selectivity filter formation in the pore region. Using site-directed mutagenesis, reduction in the molecular weight of one bulky amino acid residue in the ar/R region restricted urea and boric acid permeability. Therefore, Aqp10.2 acquired a selectivity filter during evolution, and structural differences in this selectivity filter are responsible for the variable solute permeability of aquaglyceroporins.

## Introduction

Living organisms use membrane transporters to maintain adequate concentrations of solutes, such as nutrients and toxic compounds, to ensure internal stability while adjusting to the surrounding environment. In addition to diffusion across the lipid bilayer, the transport of small molecules, such as water, occurs through channel proteins, such as aquaporins. They comprise water-specific aquaporins (wAqps), which are water-permeable, and aquaglyceroporins, which transports small uncharged solutes, such as glycerol and urea, in addition to water (1-3). Particularly, aquaglyceroporins are essential for nutrient transport and crucial for various physiological processes.

Structural analyses have revealed the mechanisms responsible for the difference in the substrate selectivity of wAqps and aquaglyceroporins (4). The Asn-Pro-Ala (NPA) motif and aromatic/Arg (ar/R) selectivity filter modulate the substrate specificity of Aqps. NPA motifs are highly conserved among Aqp family members, and two copies of the NPA motif are localized at the center of the channel pore. The ar/R motif, located on the extracellular side of the NPA motif, forms the narrowest part of the pore and is generally considered important for channel selectivity (4). wAqps form smaller pores, whereas aquaglyceroporins form larger pores, suggesting that the difference in the amino acid composition of the ar/R region influences the pore size and substrate selectivity of these proteins. Structural studies have revealed the exceptionally wide selectivity (ar/R) filter and unique cytoplasmic gate of human Aqp10 (5). These studies suggest the mechanism by which wAqps restrict the passage of large molecules, whereas aquaglyceroporins transport molecules larger than water.

Aquaglyceroporins (Aqp3, 7, 9, and 10) and water- and urea-permeable aquaporins (Aqp8) have different solute permeabilities. All aquaglyceroporins (Aqp3, 7, 9, and 10) are permeable to both glycerol and water. Additionally, Aqp7, 9, and 10 are permeable to urea, but the urea permeability of Aqp3 is much weaker or undetectable (6). Aqp8 also has a different substrate profile than other aquaglyceroporins; therefore, it is categorized in the water and urea channel subfamilies, but not in the aquaglyceroporin subfamily (7-9). However, the mechanisms underlying these differences in solute permeability have not been well-studied. A recent study on solute selectivity using ar/R-region mutants of human wAqps and aquaglyceroporins by Kitchen et al. (6) revealed that pore size alone is insufficient to explain the solute selectivity of Aqps and that substrate discrimination by Aqps depends on a complex interplay between the actual residues forming the ar/R region, physical size and chemical properties of the filter created by these residues, and the structure in which they are situated.

Previously, we reported the evolutionary specialization in the permeability of Aqp10s that can be used to investigate the mechanism of solute selectivity in aquaglyceroporins. Water, glycerol, urea, and boric acid permeabilities are plesiomorphic activities of Aqp10s that are conserved in Aqp10s of tetrapods, lobe-finned fish, and Aqp10.1 paralogs of ray-finned fish. Conversely, the permeabilities of the Aqp10.2 paralogs to urea and boric acid were much weaker than those of plesiomorphic Aqp10s (10). However, the molecular mechanism by which Aqp10.2 affects urea and boric acid transport and the details of evolutionary lineage of Aqp10s remain unclear.

This study aimed to address the molecular and structural bases underlying the specific solute permeabilities of Aqp10.1 and Aqp10.2. Phylogenetic analysis and multiple sequence alignments of the amino acid sequences of hypothetical ancestral Aqp10, tetrapod Aqp10, fish Aqp10.1, and fish Aqp10.2 sequences were performed to compare their selectivity filter domains. Three-dimensional structural models revealed the pore domain structures, which led to a novel hypothesis. Furthermore, we investigated the effects of candidate amino acid residues on the solute selectivity of Aqp10s using site-directed mutagenesis. We found that the sum of the molecular weights of the four amino acid sites in the ar/R region, if more than one is bulky, contributed to the solute selectivity of Aqp10s. These results provide structural insights into the narrowly tuned filter that is responsible for the solute selectivity of aquaglyceroporins.

## Results

### Reconstruction of ancestral Aqp10s

Differences in the solute permeability of Aqp10 between tetrapods and fish have been reported (10). However, the mechanism by which the urea and boric acid permeabilities of Aqp10.2 were lost or weakened remains unknown. Y212 (at position 3) limits the urea permeability of human Aqp3 (6). However, the African-clawed frog Aqp10 is urea-permeable despite the presence of a tyrosine residue at the same site, making it difficult to identify the specific amino acid residue affecting the solute permeability of Aqp10s. Therefore, by comparing their respective ancestral sequences as representatives rather than lone entities, we sought to determine the detailed mechanism of solute selectivity in Aqp10, Aqp10.1, and Aqp10.2. We also aimed to determine whether (i) Aqp10.2 lost the structure necessary for urea and boric acid transport or (ii) Aqp10.2 gained a filter that restricted urea and boric acid transport. To estimate which amino acid substitution occurred in the ancestor of Aqp10.2 clade, we first determined the representative sequences of Aqp10.1 and Aqp10.2. We performed ancestral sequence estimation via phylogenetic analysis using FastML (Fig. S1 and Table S2). The ancestral gene for Aqp10 diverged into two of N4, corresponding to the ancestral Aqp10 in Sarcopterygii, and N21, corresponding to the ancestral Aqp10 in ray-finned fish; N21 further diverged into two of N22, corresponding to the ancestral Aqp10.1, and N42, corresponding to the ancestral Aqp10.2. This result is consistent with the tandem duplication observed in the ancestral ray-finned fish. In addition, N42 produced N44 (via N43), corresponding to the ancestral Aqp10.2b. This represents whole-genome duplication.

To verify that the predicted sequences exhibited the expected solute selectivity, swelling assays of N22, N42, or N44 expressed in *Xenopus* oocytes was performed. N22 was used as a model sequence for Aqp10.1, N42 for Aqp10.2 before teleost-specific genome duplication (TGD), and N44 for Aqp10.2b generated from ancestral Aqp10.2 (Fig. 1A). Notably, at the upper node N21, N21-injected oocytes did not respond to the hypoosmotic solution, precluding further functional analyses (data not shown). Oocytes expressing N22, N42, or N44 showed significant volume gain and increased *P*_water_ in the hypoosmotic solution (Fig. 1B and Table S1), suggesting that these ancestral Aqp10s function as water channels in the plasma membranes of oocytes. The glycerol, urea, and boric acid permeabilities of oocytes expressing each protein were analyzed using swelling assays with an iso-osmotic solution containing 180 □ mM glycerol, urea, or boric acid, respectively. All oocytes showed significant volume gain and increase in *P*_glycerol_, suggesting the role of Aqp10s as glycerol channels. Interestingly, *P*_urea_ and *P*_boric acid_ of N42 and N44 were significantly lower than those of N22, suggesting that N22 exhibits solute selectivity as plesiomorphic Aqp10s, whereas N42 and N44 exhibit solute selectivity as Aqp10.2 (Fig. 1). The above physiological data confirmed that N22, N42, and N44 are solute permeable as expected.

**Fig. 1.**
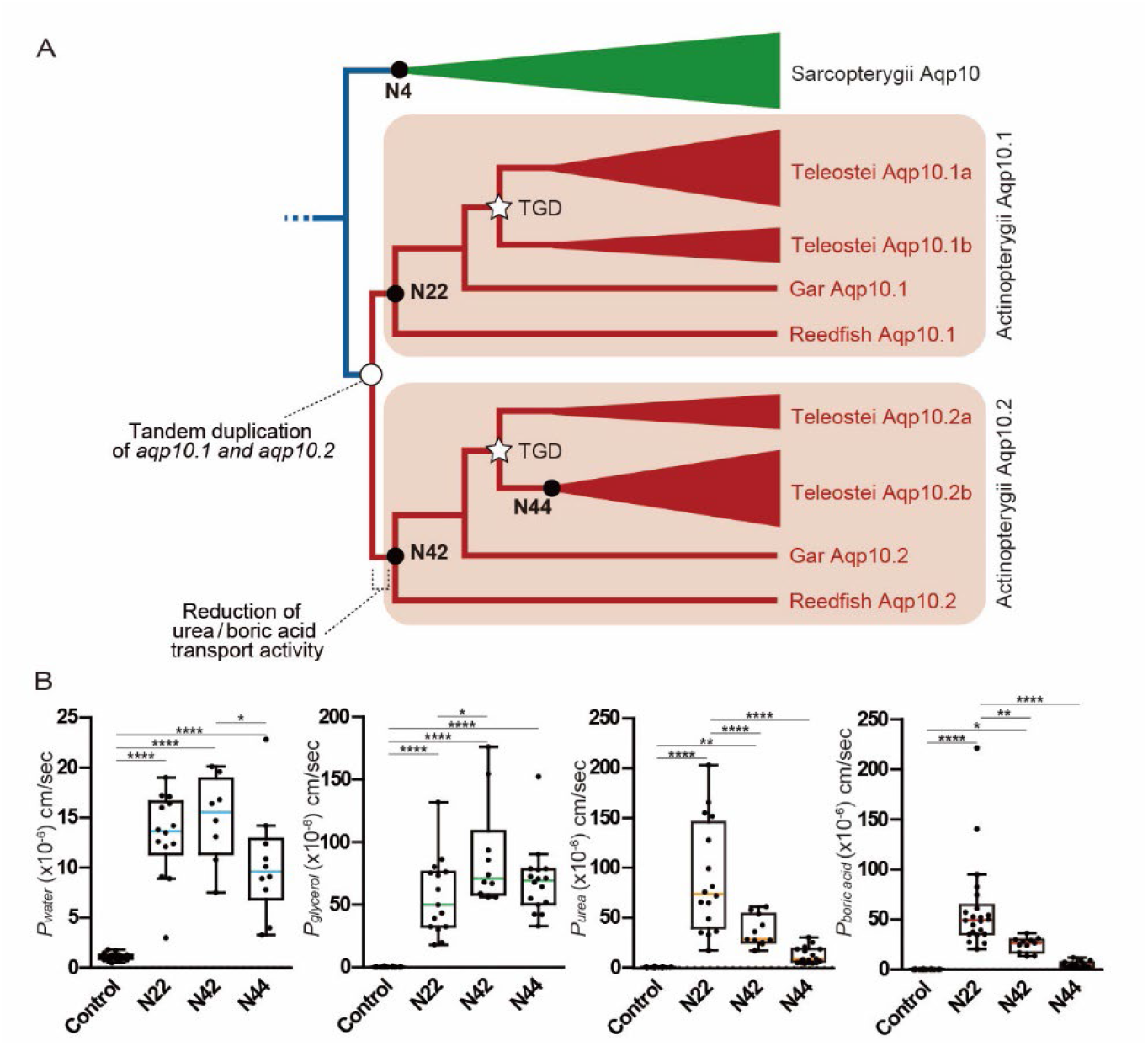
Phylogeny and water/solute permeabilities of ancestral Aqp10s. (*A*) Phylogeny of ancestral Aqp10s, especially Aqp10.1 and Aqp10.2. N22, N42, and N44 indicate the ancestral nodes of Aqp10.1, Aqp10.2, and Aqp10.2b, respectively. See Fig. S1 for details. (*B*) Solute permeabilities (volume change) of water (*P*_water_), glycerol (*P*_glycerol_), urea (*P*_urea_), and boric acid (*P*_boric acid_). Data were plotted using box-and-whisker plots (median: 25–75 percentile; 1.5 × interquartile). Statistical significance for N22, N42, and N44 was assessed via a one-way analysis of variance (ANOVA) followed by Tukey’s test (\*\*\*\**p* < 0.0001; \*\*\**p* < 0.001; \*\**p* < 0.01; \**p* < 0.05). The sample number is listed in Table S1.

### Structural models of ancestral Aqp10s and physical quantities important for their solute selectivity

To investigate the mechanism underlying solute selectivity, we conducted structural modeling of the ancestral Aqp10s N4 (Sarcopterygii), N22 (Actinopterygii, Aqp10.1), and N42 (Actinopterygii, Apq10.2) using the human Aqp10 structure as a template (Fig. 2). Unlike many ion channels, such as monomeric or dimeric, Aqp forms a tetramer with four-fold symmetry, and each of its subunits has six transmembrane helices and a pore forming four amino acid sites with the ar/R motif (positions 1–4; Fig. 2A). Previous studies on Aqps have indicated that wAqp forms smaller pores than aquaglyceroporins do, suggesting that differences in the amino acid composition of the ar/R region modulate water and substrate selectivity by affecting pore size (4). However, the substrate selectivity of aquaglyceroporins cannot be inferred solely from their amino acid residues. Therefore, we focused on the sum of molecular weights of the amino acid residues at the four ar/R sites in the model sequences. The sum of the molecular weights of the three residues centered on ar/R at the four sites was compared, considering the surrounding environment. Position 3 was scaled (x0.6) because it was on a flexible loop, whereas positions 1, 2, and 4 were not scaled because they were on a relatively rigid helix. The sum of the scaled molecular weights of the three residues in the ar/R region at positions 1–4 of N42 was greater than those of N4 and N22 (Fig. 2B). In the structural models, the estimated pore size of N42 was larger than those of N4 and N22 (Fig. 2C). These results indicate that the sum of the molecular weights of the these residues and the pore size influences the permeability of urea and boric acid.

**Fig. 2.**
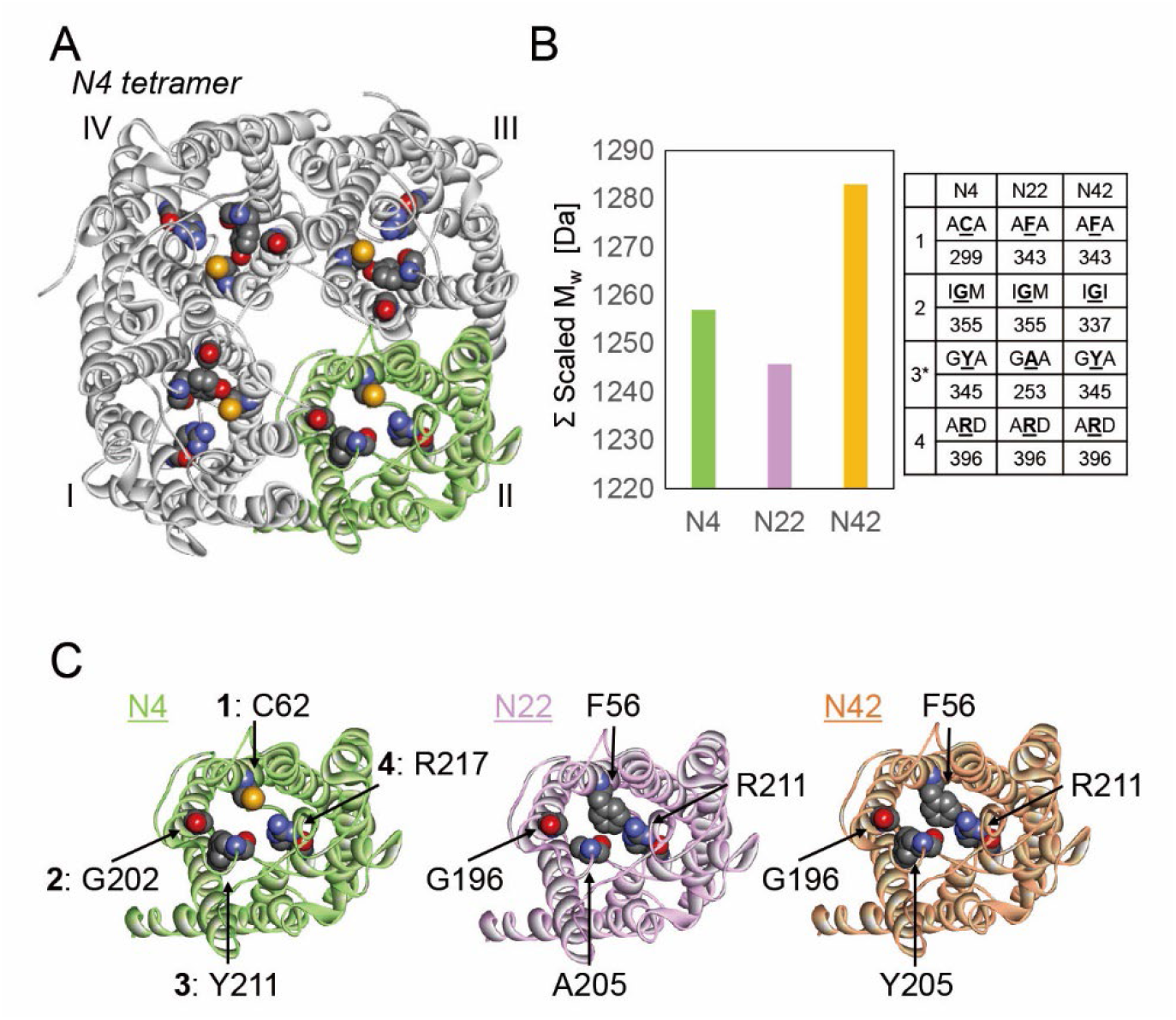
Model structures and pore molecular weights of N4, N22, and N42. (*A*) Model structure of the N4 tetramer (view from the periplasmic side). The four subunits: I (gray), II (green), III (gray), and IV (gray), and ar/R motif residues are represented in ribbon and sphere, respectively. (*B*) Sum of the scaled molecular weight (Mw) of the three residues in the ar/R region at positions 1–4 of N4, N22, and N42. Raw data at each site are shown in the right table, and the values at position 3 are scaled in the summation, due to the flexibility of its loop. (*C*) Subunit structures of N4, N22, and N42. The ar/R motif residues are shown in sphere (positions 1–4 are indicated in N4).

To determine the relationship between the sum of the molecular weights in the ar/R region and the evolutionary conservation in the actual sequences, we conducted phylogenetic analyses of 293 Aqp10 sequences (Fig. 3 and Table S3). The sum of the molecular weights of the four sites correlated well with differences in the orthologous or paralogous gene groups of tetrapods Aqp10, Aqp10.1, and Aqp10.2, and that of Aqp10.2 is larger than those of Aqp10 or Aqp10.1. Additionally, the residues at the third position of the ar/R region appeared to be G or A in Aqp10.1, and Y in Aqp10.2. These results suggest that the sum of the molecular weights, if more than one residue is bulky, determines the pore properties. This motivated us to perform further analyses using site-directed mutagenesis and the Xenopus oocyte swelling assay.

**Fig. 3.**
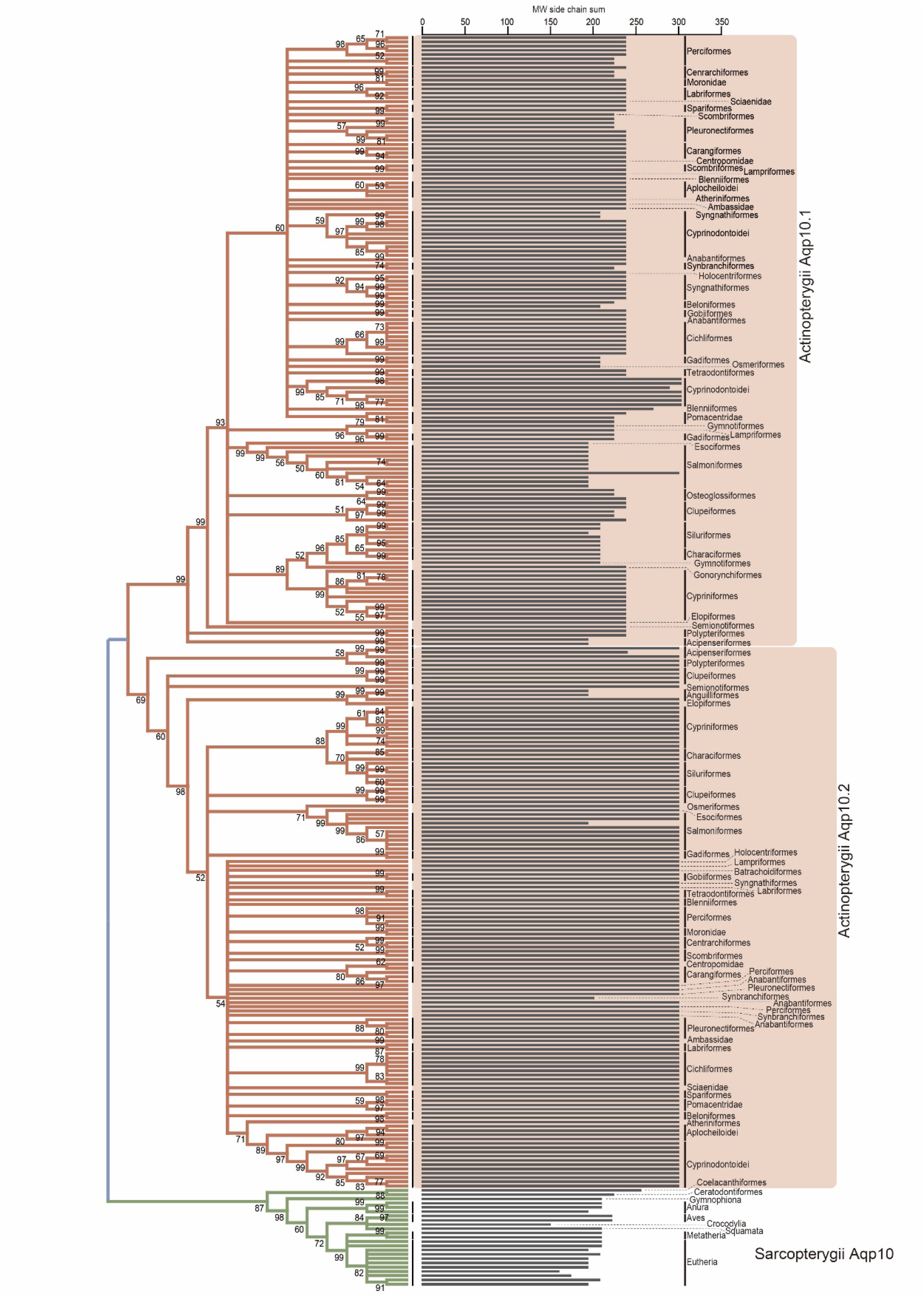
Associations between the phylogenetic tree and pore molecular weights of Aqp10, Aqp10.1/1a, and Aqp10.2/2b. Left, phylogenetic tree of Aqp10, Aqp10.1/1a, and Aqp10.2/2b. Right, sum of molecular weights in the ar/R region of Aqp10.

### Effects of site-directed mutagenesis on the selectivity filter of N42 (ancestral Aqp10.2) for urea and boric acid transport

We examined whether mutations in the N42 residues would cause changes in urea and boric acid permeability as a result of pore size changes. Using the *Xenopus* oocyte swelling assay, we generated and analyzed two mutations, F56G (position 1) and Y205A (position 3), in N42 using the *Xenopus* oocyte swelling assay, both of which reduced the sum of the molecular weights of the four amino acid sites in the ar/R region (Fig. 4A). Oocytes expressing N42 mutants showed significant volume gains and increases in *P*_*water*_ in the hypoosmotic solution (Fig. 4A and Table S1), suggesting that these N42 mutants act as water channels in the plasma membranes of the oocytes. Moreover, in an iso-osmotic solution containing glycerol, oocytes expressing N42 mutants showed significant volume gains and increased *P*_*glycerol*_ (Fig. 4A), suggesting that they also act as glycerol channels. In these experiments, the average values of *P*_*glycerol*_ for N42 F56G and N42 Y205A were slightly 0.70- and 0.77-times lower than those of N42, respectively, whereas the averages of *P*_*water*_ were not significantly different between the N42 WT and mutants. Furthermore, in an iso-osmotic solution containing urea and boric acid, oocytes expressing N42 mutants showed significant cell volume gains and increases in *P*_*urea*_ and *P*_*boric acid*_ (Fig. 4A), suggesting that N42 mutants act as urea and boric acid channels. The average values of *P*_*urea*_ and *P*_*boric acid*_ of N42 F56G and N42 Y205A were 2.7–3.5 times higher than those of N42. These results provide evidence that the sum of the scaled molecular weights of ar/R alters *P*_*urea*_ and *P*_*boric acid*_.

**Fig. 4.**
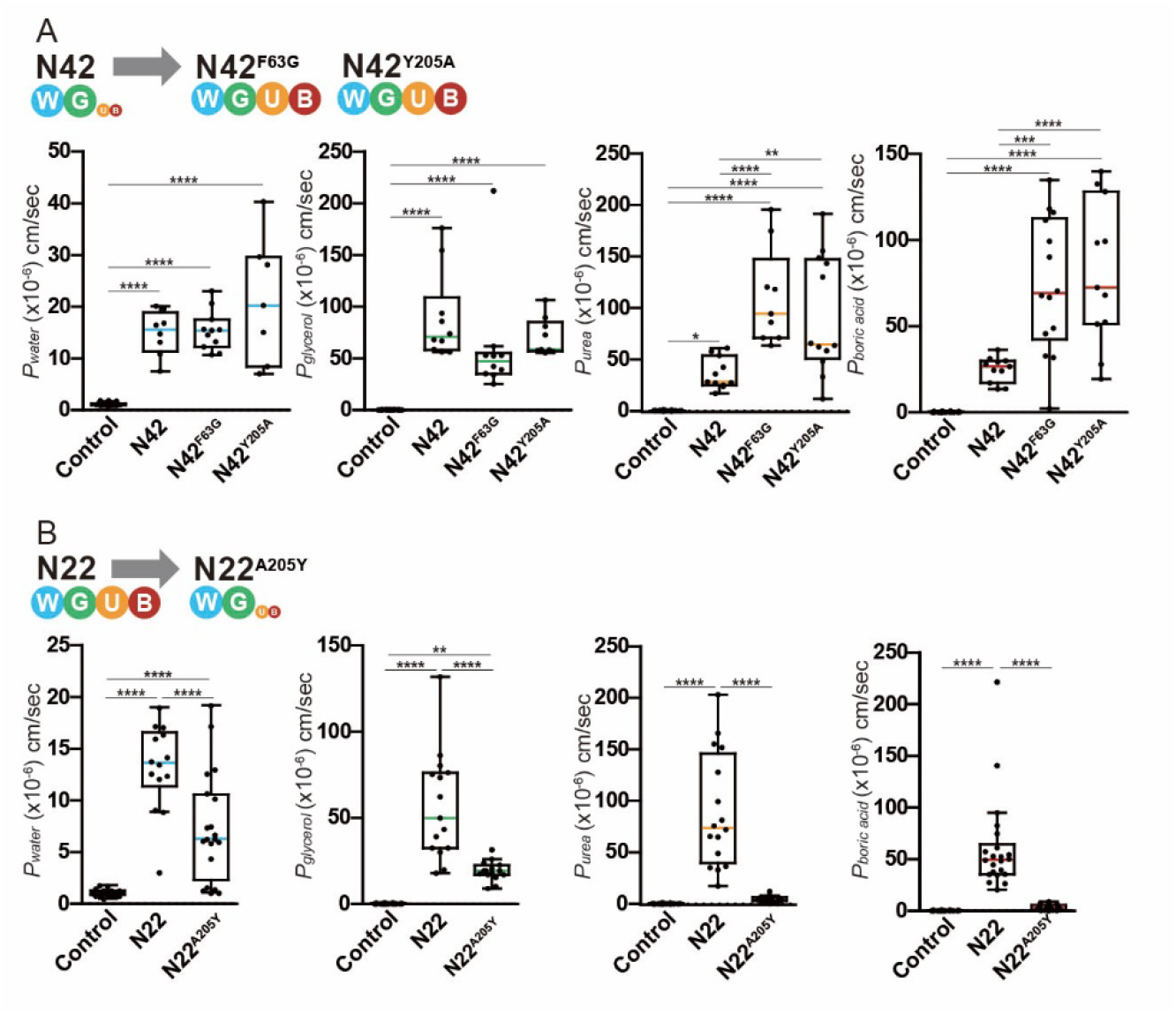
Water and solute permeabilities of N42, F56G and Y205A (N42 mutants), N22, and A205Y (N22 mutant). (*A*) Water (*P*_water_), glycerol (*P*_glycerol_), urea (*P*_urea_), and boric acid (*P*_boric acid_) permeabilities (volume changes) in oocytes expressing N42 and its mutant compared with those in control oocytes. Data were plotted using box-and-whisker plots (median: 25–75 percentile; 1.5 × interquartile). Statistical significance was assessed via a one-way analysis of variance (ANOVA) followed by Tukey’s test (\*\*\*\**p* < 0.0001; \*\*\**p* < 0.001; \*\**p* < 0.01). Samples are listed in Table S1. (*B*) Water (*P*_water_), glycerol (*P*_glycerol_), urea (*P*_urea_), and boric acid (*P*_boric acid_) permeabilities in oocytes expressing N22 and its mutant compared with those in control oocytes. Data were plotted using box-and-whisker plots (median: 25–75 percentile; 1.5 × interquartile). Statistical significance was assessed via a one-way analysis of variance (ANOVA) followed by Tukey’s test (\*\*\*\**p* < 0.0001; \*\**p* < 0.01). The sample number is listed in Table S1.

### F56G or Y205A mutations in Aqp10.2 of real “ancient” ray-finned fish for urea and boric acid permeability

We next investigated whether mutations in the corresponding residues in the gray bichir Aqp10.2 (PseAqp10.2), would cause similar changes in selectivity filter function. We previously showed that the permeability of the PseAqp10.2 to urea and boric acid was much weaker than that of PseAqp10.1 in the preceding study (10). Here, we generated two mutations (the same as above) in PseAqp10.2, which reduced the sum of the molecular weights of the four amino acid sites in the ar/R region and analyzed them using a *Xenopus* oocyte swelling assay (Fig. 5A). As expected, oocytes expressing PseAqp10.2 mutants showed significant volume gain and increases in both *P*_water_ (hypoosmotic solution) and *P*_glycerol_ (iso-osmotic solution) (Fig. 5A and Table S1), suggesting that these PseAqp10.2 mutants act as water and glycerol channels in the plasma membranes of oocytes. In addition, in an iso-osmotic solution containing urea and boric acid, oocytes expressing PseAqp10.2, but not wild-type PseAqp10.2, showed significant cell volume gains and increases in *P*_urea_ and *P*_boric acid_ (Fig. 5A), suggesting that PseAqp10.2 mutants have acquired permeability to urea and boric acid. The average values of *P*_urea_ and *P*_boric acid_ of PseAqp10.2^F56G^ and PseAqp10.2^Y205A^ were 20–26-times higher than those of wild-type PseAqp10.2. These results are in good agreement with those of the N42 mutants described above and confirm the influence of the molecular weight of the pore on solute permeability.

**Fig. 5.**
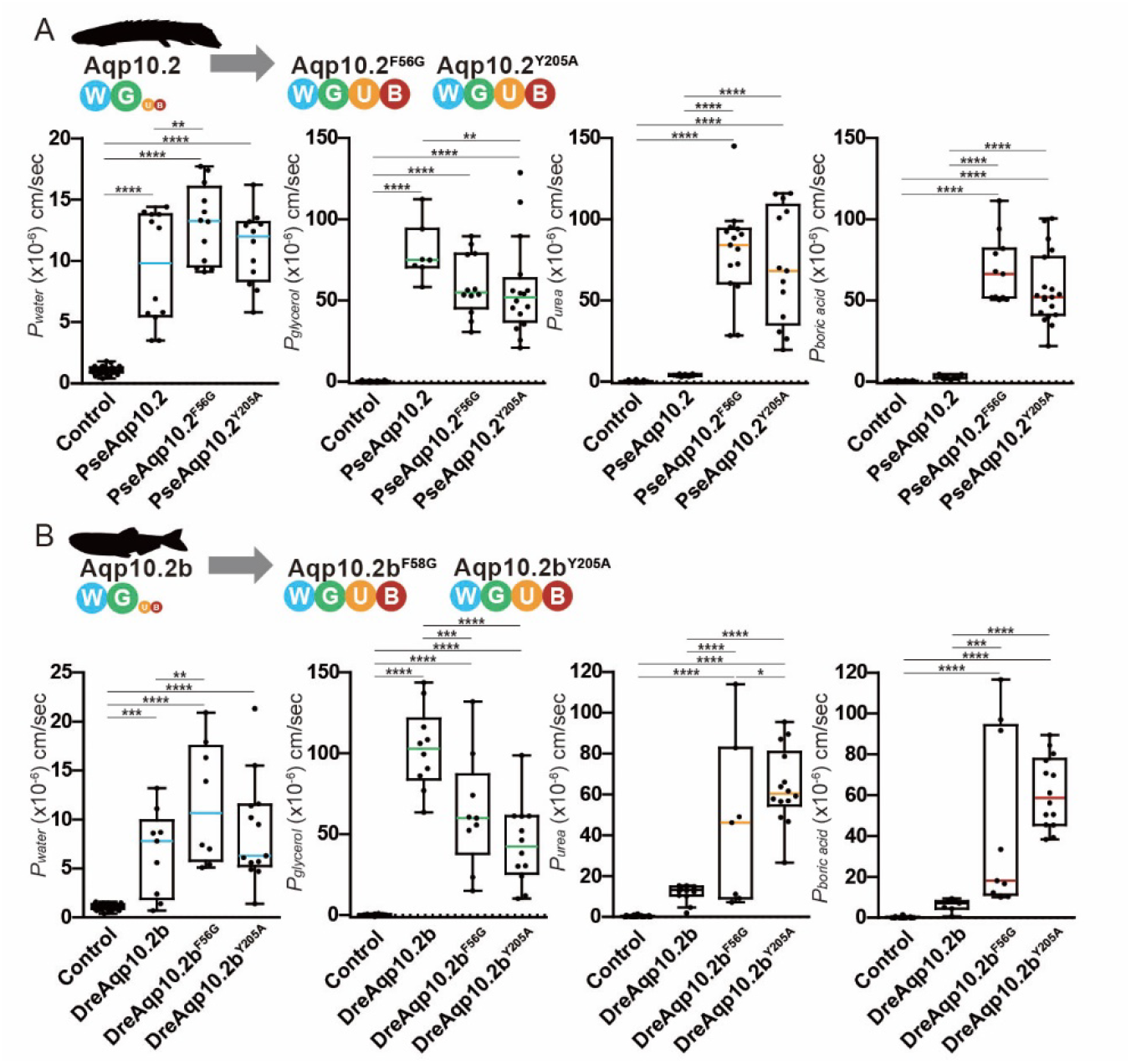
Water and solute permeabilities of Aqp10.2/2b and its mutants in gray bichirs (PseAqp10.2) and zebrafish (DreAqp10.2b). Water (*P*_water_), glycerol (*P*_glycerol_), urea (*P*_urea_), and boric acid (*P*_boric acid_) permeabilities (volume changes) of PseAqp10.2 and its mutants (*A*) and DreAqp10.2b and its mutant (*B*). Data were plotted using box-and-whisker plots (median: 25–75 percentile; 1.5 × interquartile). Statistical significance was assessed via a one-way analysis of variance (ANOVA) followed by Tukey’s test (\*\*\*\**p* < 0.0001; \*\*\**p* < 0.001; \*\**p* < 0.01; \**p* < 0.05). The sample number is listed in Table S1.

### F56G or Y205A mutations in Aqp10.2b of teleost species with two *aqp10* paralogs for urea and boric acid permeability

Gray bichir Aqp10s sequences were obtained from the genome database and artificially synthesized. To examine the sequences were cloned from actual fish, we tested whether mutations in the corresponding residues of zebrafish (DreAqp10.2b) cause similar changes in selectivity filter function. Notably, TGD generated *aqp10*.*2a* and *aqp10*.*2b* from *aqp10*.*2*, whereas zebrafish lost *aqp10*.*2a* and only had *aqp10*.*2b*. Similarly, we generated two mutations in DreAqp10.2b that reduced the sum of the molecular weights of the four amino acid sites in the ar/R region and analyzed them using the *Xenopus* oocyte swelling assay (Fig. 5B and Table S1). Oocytes expressing DreAqp10.2b mutants also showed significant volume gains and increases in *P*_water_ and *P*_glycerol_ (Fig. 5B), suggesting that these DreAqp10.2b mutants act as water and glycerol channels in the plasma membranes of oocytes. Additionally, in an iso-osmotic solution containing urea and boric acid, only oocytes expressing DreAqp10.2b mutants, but not wild-type DreAqp10.2b, showed significant cell volume gains and increases in *P*_urea_ and *P*_boric acid_ (Fig. 4B), suggesting that DreAqp10.2b mutants act as urea and boric acid channels. The average values of *P*_urea_ and *P*_boric acid_ of DreAqp10.2b mutants were 4.2–9.8 times higher than those of the wild-type DreAqp10.2b. These results were also fairly consistent with those of N42 and PseAqp10.2 mutants as above, although there was a slight difference in the sequence assembly between artificially synthesized sequences and cloned sequences. Taken together, the reduction in the sum of the molecular weights of residues in the ar/R region resulted in increased permeability to urea and boric acid compared with that of the wild-type in all experiments.

### Effects of site-directed mutagenesis on the generation of N22 and Aqp10.1/1a selectivity filters for urea and boric acid transport

In the above experiments, a selectivity filter for Aqp10.2/2b was successfully modified using site-directed mutagenesis. Next, we examined whether mutations in the residues of plesiomorphic Aqp10s cause a decrease in urea and boric acid permeabilities due to pore size changes. First, we examined a mutant of N22, A205Y, using the *Xenopus* oocyte swelling assay (Fig. 4). Oocytes expressing the N22 mutant showed significant volume gain and increase in both *P*_*water*_ and *P*_*glycerol*_ (Fig. 4B and Table S1), suggesting that the N22 mutant acts as a water and glycerol channel in the plasma membranes of oocytes. In these experiments, the average values of *P*_*water*_ and *P*_*glycerol*_ for N22 A205Y were 0.55- and 0.34-times lower than those of N22, respectively. In the iso-osmotic solution containing urea or boric acid, N22, but not the N22 A205Y mutant, showed significant volume gain and increase in *P*_*urea*_ and *P*_*boric acid*_ (Fig. 4B), suggesting that N22 A205Y does not act as a urea or boric acid channel.

Next, we evaluated the PseAqp10.1 and DreAqp10.1a mutants using *Xenopus* oocyte swelling assays (Fig. S2). PseAqp10.1 A205Y and DreAqp10.1a A201Y (position 3) mutants did not transport water, precluding further functional analysis (Figs. S2A and S2B). PseAqp10.1 S196G (position 2) mutant had little or no effect on permeability. These results indicated that urea and boric acid permeabilities were mostly transferable (from Aqp10.2 to plesiomorphic Aqp10s), whereas the formation of a selectivity filter (from plesiomorphic Aqp10s to Aqp10.2) could not be fully explained, probably due to the unknown effects of residue modifications. These findings suggest that urea and boric acid filters, which require additional amino acid residues, structural scaffolds, or machinery, were simultaneously generated during the evolution of Aqp10.

## Discussion

In this study, we propose that the presence of two or more bulky amino acids in the ar/R region, which increase the sum of the molecular weights of the ar/R region, is essential for the formation of a filter that limits urea and boric acid transport. Previous comparative analyses of aquaglyceroporins have mainly been based on comparisons of aquaglyceroporins from different organisms; however, our detailed analysis of evolutionary changes allowed us to compare the putative ancestral sequences of each paralog. As a result, the amino acid sites affecting activity were identified. Focusing on the molecular weight rather than the type of amino acid in the amino acid moiety, we compared Aqp10.2 sequences with those of Aqp10s (approximately 300 in total), and examined the effect of candidate amino acid residues on solute selectivity using a site-directed mutagenesis method. These results suggest that Aqp10.2 resulting from tandem duplication has not lost its transport ability, but has gained a filter with substrate selectivity.

Previous studies using human Aqp3 mutants, which transport water and glycerol but have low urea transport activity, have reported that it is necessary to replace Y at position 3 with A to obtain high urea permeability (6). Therefore, we initially hypothesized that Y at position 3 of ar/R might be important for regulating the urea and boric acid transport activities of Aqp10, Aqp10.1, and Aqp10.2. However, this site is Y/I in tetrapod Aqp10s, such as those in African clawed frogs and humans, which transport urea and boric acid. This suggests that the bulky Y at position 3 of ar/R was insufficient to disrupt urea transport in tetrapod Aqp10s. We then attempted to narrow the pores and construct a selectivity filter by introducing a mutation into tetrapod Aqp10s that replaced the small amino acid residue (G) at position 1 with a bulky amino acid residue (F) (Fig. S3). This mutation unexpectedly restricted the transport of glycerol, urea, and boric acid. On the other hand, even Aqp10.2/2b, with intact Y in position 3 and a mutation from F to G at position 1, transported urea and boric acid similar to tetrapod Aqp10s and ray-finned fish Aqp10.1s (Fig. 5). Aqp10.2/2b, which has a mutation from Y to A at position 3, also transports urea and boric acid (Fig. 5). This indicates that Y at position 3 alone is not sufficient, but the presence of at least two bulky amino acid residues at positions 1 and 3, which increase the sum of the molecular weights of the ar/R region, is important. This is the second report describing the substrate selectivity of urea for aquaglyceroporins, following the previous study (10). While supporting the previous report, this study extends it with a new hypothesis.

In our site-directed mutagenesis analysis, experiments that reduced the sum of the molecular weights of the ar/R region increased the permeability to urea and boric acid. However, increasing the sum of the molecular weights of the ar/R region did not always limit the permeability of urea and boric acid favorably. Thus, fewer functional constraints are required for the formation of a filter with low selectivity, and higher functional constraints are required for the formation of a filter with high selectivity. These results are consistent with an Idea proposed by Kitchen et al. (6) that pore size alone is insufficient to explain the solute selectivity of Aqps and that substrate discrimination in Aqps depends on a complex interplay between the actual residues that form the ar/R region, the physical size and chemical properties of the filter created by these residues, and the structural context in which they are situated.

Furthermore, we previously reported that Japanese pufferfish (*Takifugu rubripes*) Aqp9a transports water, urea, and boric acid, but little glycerol (11), and that the set of amino acids in ar/R was consistent with the tetrapod Aqp10 mutants examined in this study. Additionally, human Aqp8, which has similar transport substrate selectivity, has a residue H at position 1 (12), suggesting that not only the size/molecular weight of the amino acid residues but also their polarity are involved in glycerol transport. Further studies on glycerol selectivity of Aqp10s are required. Throughout the history of evolution, animals have acquired various Aqps through gene loss and duplication. In this study, we found that Aqp10.2/2b weakened the permeability of slightly larger substrates (urea and boric acid) by replacing residues in the pore region with residues of larger molecular weights. Changes in Aqps in animals, such as fish, may be due to the environmental factors in which they survive.

In conclusion, this study demonstrated that the selectivity filter and solute selectivity of aquaglyceroporins for small molecules are regulated by the amino acid residues in the ar/R region and their molecular weights, that is, simple physical quantities. Similar to the structural comparison of conventional aquaporins and aquaglyceroporins, the pore size of Aqp10.2/2b was narrower than that of Aqp10 or Aqp10.1 and acted as a barrier against urea and boric acid transport. Many aquaglyceroporins are not similar in their amino acid sequences, making it difficult to investigate their amino acids. Here, our findings indicate that aquaglyceroporins, such as Aqp10s, TruAqp9a, and HsaAqp8, have a common permeability mechanism for various external small molecules. Although the mechanism by which Aqp10s, especially Aqp10.2/2b, alter their substrate selectivity during evolution and its physiological significance remain unknown, we succeeded in partially reconstituting the Aqp10 filter by modifying the selectivity filter of Aqp10.2/2b in an in vitro swelling assay in this study. Therefore, our study provides a better understanding of the contribution of Aqps to the small-molecule absorption of epithelial cells and their regulation from structural and evolutionary perspectives.

## Materials and Methods

### Ancestral sequence estimation based on phylogenetic analysis

Sequences of Aqp10 and outgroup Aqp9 were obtained from the study of Yilmaz et al. (13), and Aqp10 sequences of herring and shad were added to this dataset. The amino acid sequences were aligned using MAFFT version 7 with the linsi option and used to construct a maximum-likelihood tree with IQ-TREE v1.6.12 (-m MFP) (14). Ancestral sequences were reconstructed using the maximum likelihood method with FastML v3.1 using the JTT + G model (15).

### Phylogenetic analysis

Evolutionary history was inferred using the maximum likelihood method with the JTT matrix-based model (16). The tree with the highest log likelihood (–53707.71) is shown (Fig. 3). The percentages of trees in which the associated taxa were clustered together are shown below the branches. The initial tree(s) for the heuristic search were obtained by applying the neighbor-joining method to a matrix of pairwise distances estimated using the JTT model. The analysis included 293 amino acids. A total of 760 positions were involved in the final dataset. Evolutionary analyses were conducted using MEGA11 (17). Here, we constructed a condensed tree. If several branches have bootstrap values < 50, a condensed tree with 50 bootstrap values will have a multifurcating tree, with all its branch lengths reduced to zero.

### Structural modeling of ancestral Aqp10s (N4, N22, and N42)

Model structures of the predicted ancestral sequences N4, N22, and N42 (Table S2) were constructed using the human Aqp10 structure (PDB 6F7H (5) as a template using the SWISS-MODEL server (18). The template sequence had a sequence identity of 63.23, 56.92, and 63.18% and coverage of 0.88, 0.75 and 0.77 with the N4, N22, and N42 sequences, respectively. As a high sequence identity (> 60%) and wide coverage (> 0.75) were obtained for all three, we adopted the SWISS-MODEL model structures instead of AlphaFold prediction (19).

### Cloning of Aqp10s using the human, African clawed frog, gray bichir, zebrafish, and ancestral sequences

All animal protocols and procedures were conducted in accordance with the National Institutes of Health Guide for the Care and Use of Laboratory Animals and a manual approved by the Institutional Animal Experiment Committee of the Tokyo Institute of Technology. Full-length cDNAs of Aqp10s from human, African-clawed frogs, and zebrafish were isolated from previously prepared intestinal cDNA of each species (20, 21). Full-length cDNAs of Aqp10s from gray bichir Aqp10s and ancestral sequences were chemically synthesized (Eurofins Genomics, Tokyo, Japan), as previously described (10). cDNAs were amplified via polymerase chain reaction (PCR) using a high-fidelity DNA polymerase (KOD One PCR Master Mix; Toyobo, Osaka, Japan), and the primers were designed based on the genomic database (Table S4). cDNAs were subcloned into pGEMHE (22) using the In-Fusion Snap Assembly Master Mix (Takara Bio, Shiga, Japan) with gene-specific primers and 15-bp sequences complementary to the ends of the linearized pGEMHE (Table S4) and sequenced.

### Site-directed mutagenesis targeting the Aqp10 pore region

Point mutations were introduced into Aqp10s using a reaction mixture containing high-fidelity DNA polymerase (KOD One PCR Master Mix; Toyobo), primers (Table S4), and pGEMHE-containing *aqp10* as a template via PCR. Full-length cDNAs of mutated DreAqp10.2 were chemically synthesized (Eurofins Genomics) and subcloned as described above, and all mutation sites were confirmed via DNA sequencing.

### Expression of Aqp10s in *Xenopus* oocytes

The plasmids were linearized with NotI (Takara Bio), and capped RNAs (cRNAs) were transcribed *in vitro* using atheT7 Message Machine kit (Thermo Fisher Scientific, Waltham, MA, USA). *Xenopus laevis* oocytes were dissociated with collagenase, as previously described (11, 12, 23), and injected with 50 nL of water or a solution containing 0.5 ng/nL cRNA (25 ng/oocyte) using the Nanoject-II injector (Drummond Scientific, Broomall, PA, USA). Oocytes were incubated in the OR3 medium at 16 °C and observed for 5 d after injection. OR3 medium (1 L) consisted of 0.7% w/v powdered Leibovitz L-15 medium with L-glutamine (Thermo Fisher Scientific), 50 mL of 10,000 U penicillin and 10,000 U streptomycin solution in 0.9% NaCl (Nacalai Tesque Inc., Kyoto, Japan), and 5 mM HEPES (pH 7.50). The osmolality was adjusted to 200 mOsmol/kg with NaCl powder (11, 12, 23). The frogs were handled, and the oocytes were harvested as described in the “Cloning of Aqp10s using the human, African clawed frog, gray bichir, zebrafish, and ancestral sequences” section.

### Swelling assay

Oocyte swelling was monitored using a stereomicroscope (SZX9; Olympus, Tokyo, Japan) equipped with a charge-coupled device camera (DS-Fi2; Nikon, Tokyo, Japan), as previously described (11, 12). The oocyte volume was calculated assuming a spherical geometry. Oocytes incubated with ND96 (approximately 200 mOsmol/kg) were transferred to two-fold diluted ND96 (approximately 100 mOsmol/kg) for water transport assays. For the glycerol, urea, and boric acid transport assays, oocytes were transferred to an isotonic solution containing ND96 supplemented with 180 mM glycerol, urea, or boric acid instead of NaCl and adjusted to an osmolality of approximately 200 mOsmol/kg. Water permeability (*P*_water_) was calculated from the osmotic swelling data and molar volume of water (*V*_*w*_ = 18□cm^3^/mol) as follows (24): *P*_water_ = [*V*_*o*_□× □*d*(*V*/*V*_*o*_)/*dt*]/[*S□*× □*V*_*w*_□× □ (osm_in_ – osm_out_)], where *S* is the initial oocyte surface area. Solute permeability (*P*_solute_) was calculated from the swelling data, total osmolality of the system (osm_total_ = 200 mOsmol/kg), and solute gradient (sol_out_ – sol_in_) as follows (25): *P*_solute_ = osm_total_□× □ [*V*_*o*_□× □*d*(*V*/*V*_*o*_)/*dt*]/[*S* × (sol_out_ – sol_in_)]. The water, glycerol, urea, and boric acid transport activities of each Aqp10 strain were evaluated using oocytes from the same animal, and the experiment was repeated in at least three frogs. Quantitative data are represented as the mean ± standard deviation in Table S1. *P*_*water*_ and *P*_*solute*_ values were compared between the Aqp10-expressing and control oocytes, and the statistical significance was assessed using one-way analysis of variance, followed by Tukey’s test with GraphPad Prism software (version 8; GraphPad, San Diego, CA, USA) to visualize the results in box plots.

## Acknowledgments

We would like to thank Shin-ichiro Hidaka, Natsue Yamamoto, Michiko Kotani, Tomoko Narisawa, Sachiko Muto, Satomi Asano, Yoko Yamamoto, Nana Shinohara, Atsumi Sakaguchi, the Biomaterials Analysis Division, and Open Research Facilities for Life Science and Technology at the Tokyo Institute of Technology for their technical assistance in this study. This study was financially supported by the Japan Society for the Promotion of Science (JSPS) KAKENHI (grant numbers: 17H03870, 21H02281, 21K14781, and 19H03272).

## Data Availability

All study data are included in the article and/or SI Appendix.

