## Supplementary material for "Selectivity and evolution of Aqp10 in solute permeability influenced by pore molecular weight": Table S1

**Table S1.** Water and solute permeabilities of Aqp10s in oocytes.

| Protein | $P_{water}$ (× $10^{-6}$ cm/s, 100 mosM inside osmotic gradient) | $P_{glycerol}$ (× $10^{-6}$ cm/s, 180 mM outside solute gradient) | $P_{urea}$ (× $10^{-6}$ cm/s, 180 mM outside solute gradient) | $P_{boric\ acid}$ (× $10^{-6}$ cm/s, 180 mM outside solute gradient) | Figure # |
|---|---|---|---|---|---|
| Control | 1.1 ± 0.3(24) | 0.2 ± 0.2 (24) | 0.09 ± 0.29 (24) | 0.35 ± 0.23 (24) | Figure. 1 |
| N22 | 13.2 ± 4.0 (14) | 56.7 ± 29.9 (15) | 89.5 ± 53.6 (16) | 60.9 ± 43.6 (22) | |
| N42 | 14.9 ± 4.0 (8) | 89.0 ± 40.2 (10) | 36.3 ±14.4 (11) | 24.9 ± 7.0 (11) | |
| N44 | 10.4 ± 5.2 (10) | 68.6 ± 26.6 (16) | 12.7 ± 8.1 (15) | 5.1 ± 2.9 (16) | |
| Control | 1.2 ± 0.3 (23) | 0.2 ± 0.2 (23) | 0.1 ± 0.3 (22) | 0.3 ± 0.2 (24) | Figure. 4 |
| N42 (Figure 1) | 14.9 ± 4.0 (8) | 89.0 ± 40.2 (10) | 36.3 ±14.4 (11) | 24.9 ± 7.0 (11) | |
| N42 F56G | 15.4 ± 3.7 (11) | 60.8 ± 51.5 (10) | 110.5 ± 44.4 (9) | 74.0 ± 37.7 (14) | |
| N42 Y205A | 21.2 ± 19.7 (7) | 70.7 ± 17.1 (9) | 92.6 ± 55.1 (12) | 80.9 ± 39.9 (11) | |
| Control | 1.1 ± 0.3 (34) | 0.1 ± 0.3 (34) | 0.1 ± 0.4 (33) | 0.2 ± 0.2 (40) | Figure. 5A |
| PseAqp10.2 | 9.4 ± 4.4 (12) | 79.6 ± 16.6 (7) | 4.0 ± 0.6 (8) | 2.8 ± 1.2 (7) | |
| PseAqp10.2 F56G | 13.0 ± 3.0 (12) | 59.5 ± 18.3(12) | 79.4 ± 27.9 (15) | 68.8 ±19.7(11) | |
| PseAqp10.2 Y205A | 11.1 ± 2.9 (12) | 57.0 ± 28.6 (16) | 70.9 ± 34.4 (13) | 57.1 ± 21.3 (19) | |
| Control | 1.1 ± 0.3 (32) | 0.2 ± 0.3 (37) | 0.3 ± 0.3 (36) | 0.3 ± 0.3 (33) | Figure. 5B |
| DreAqp10.2b | 6.6 ± 4.1 (9) | 102.8 ± 23.9 (10) | 12.0 ± 3.9 (14) | 6.1 ± 2.7 (7) | |
| DreAqp10.2b F56G | 12.4 ± 5.8 (8) | 63.6 ± 33.8 (9) | 45.6 ± 37.9 (7) | 45.2 ± 41.0(9) | |
| DreAqp10.2b Y205A | 8.8 ± 5.1 (13) | 43.7 ± 24.0 (12) | 63.9 ± 17.9 (14) | 61.3 ± 16.6 (14) | |
| Control | 1.1 ± 0.3 (24) | 0.2 ± 0.2 (24) | 0.09 ± 0.3 (24) | 0.3 ± 0.2 (24) | Figure. 4 |
| N22 (Figure 1) | 13.2 ± 4.0 (14) | 56.7 ± 29.9 (15) | 89.5 ± 53.6 (16) | 60.9 ± 43.6 (22) | |
| N22 A205Y | 7.3 ± 5.1 (20) | 19.1 ± 5.5 (15) | 4.6 ± 2.9 (18) | 3.6 ± 2.5 (20) | |
| Control | 1.1 ± 0.3 (35) | 0.2 ± 0.3 (35) | 0.2 ± 0.5 (35) | 0.2 ± 0.2 (40) | Figure. S2A |
| PseAqp10.1 | 10.0 ± 2.2 (10) | 73.2 ± 30.9 (9) | 85.2 ± 23.4 (7) | 61.1 ± 25.7 (7) | |
| PseAqp10.1 S196G | 11.5 ± 1.4 (10) | 52.7 ± 21.5 (13) | 70.6 ± 26.6 (13) | 74.6± 24.9 (15) | |
| PseAqp10.1 A205Y | 1.5 ± 0.3 (13) | 1.7 ± 1.0 (12) | 1.9 ± 1.4 (12) | 2.4 ± 0.7 (13) | |
| PseAqp10.1 S196G_A205Y | 1.5 ± 0.2 (12) | 1.5 ± 0.6 (12) | 0.5 ± 0.4 (12) | 0.6 ± 0.2 (12) | |
| Control | 1.0 ± 0.3 (20) | 0.1 ± 0.3 (25) | 0.2 ± 0.2 (24) | 0.3 ± 0.3 (21) | Figure. S2B |
| DreAqp10.1a | 12.3 ± 5.3 (9) | 45.9 ± 25.8 (17) | 79.6 ± 27.3 (11) | 48.0 ± 21.2 (5) | |
| DreAqp10.1a A201Y | 0.8 ± 0.4 (12) | 0.4 ± 0.4 (12) | 1.1 ± 0.7 (12) | 1.1 ± 0.9 (11) | |
| Control | 1.1 ± 0.4 (38) | 0.5 ± 0.7 (22) | 0.8 ± 1.0 (34) | 0.5 ± 0.5 (36) | Figure. S3A |
| HsaAqp10 | 3.3 ± 2.1 (30) | 9.7 ± 8.4 (15) | 10.7 ±5.4 (22) | 7.6 ± 6.5 (22) | |
| HsaAqp10 G62F | 4.1 ± 2.3 (10) | 0.5 ± 0.6 (12) | 3.3 ± 2.6 (13) | 5.0 ± 3.0 (10) | |
| Control | 1.2 ± 0.2 (23) | 0.2 ± 0.2 (23) | 0.4 ± 0.4 (22) | 0.2 ± 0.2 (24) | Figure. S3B |
| XlaAqp10 | 10.7 ± 2.8 (12) | 54.0 ± 22.7 (10) | 73.0 ± 13.6 (9) | 57.1 ± 20.0 (11) | |
| XlaAqp10 G86F | 6.6 ± 1.6 (13) | 0.7 ± 0.2 (12) | 19.4 ± 4.9 (12) | 11.0 ± 5.6 (15) | |

Values are represented as the mean ± standard deviation. Number in parentheses indicates the total number of oocytes assayed.
