## Supplementary material for "Selectivity and evolution of Aqp10 in solute permeability influenced by pore molecular weight": Table S2

**Table S2.** Ancestral sequences of Aqp10s.

| Name | Amino acid sequence |
|---|---|
| N4 | MGRAPSMERVRALLRIRNPLVRECLAEFLGTFVLILFGLGATAQVVTSHETKGDYLSINLA**C**ALGVTFGIYVSGGVSGA<br>HLNPAVSLSMCLLGRFPWWKLPFYVLFQILGAFLAAATVYALYYDAIQNYSGGNLTVTGPRETASIFATYPAPYLSIRN<br>GFIDQVIGTAMLLVCILAIVDSRNSPVPKGLEPVLVGLVVLTI**G**MSMGSNCG**Y**AINPA**R**DLGPRLFTYVAGWGPEVFTA<br>GNNWWWVPVVAPLVGAVLGTLLYLLFVEFHHPDPESEPQESATAKNKQKAEKGTPGTEKKKAVEVPTFKSNPRGEGSLE<br>REGEGEESLAHRL |
| N22 | MEKLRRLLRLKNRLLRECLAEFLGTFVLILFGCAAAAQVKTSHETKGQYLSINLA**F**AVGVIFAMYISRGVSGAHLNPAV<br>SLSFCVLGRFPWRKLPFYILFQILGAFLASATVFAQYYDAIMNYSGGNLTVTGPNETASIFATYPSEYLSLSNAFVDQV<br>IGTAALLLCILPLDDSRNSPAPKGLEPVLVGFVVLGI**G**MSMSSNCG**A**AINPA**R**DLGPRLFTLAAGWGTEVFTAFNNWWW<br>IPVVAPMLGGVLGACIYLLFIEFHHPDPESDSQESSTAENKTSAEAVSPGTNAKDAVSLPTSKSNPRGDGSLKESTKLH<br>IIKNPFLEGQSENSDWERESEGEECIAHRL |
| N42 | MEKVRRLLRVRNQLVRECLAEFLGTYVLILFGCGAVAQVTTSHETKGQYLSINLA**F**ALGVTFGVYVSRGVSGAHLNPAV<br>SLSMCLLGRFPWRKLPFYVLFQILGAFLAAATVYAQYYDAIMNYSGGNLTVTGPNATASIFATYPAPYLSISNGFVDQV<br>IGTAALLLCVLALGDSRNSPAPKGLEPVLVGLVVLVI**G**ISMGSNCG**Y**AINPA**R**DLGPRLFTYVAGWGTEVFTAGNNWWW<br>VPLVAPMVGALLGTAIYLLFIELHHPAPESESQDSATGKNKYAVELEEGKSNPRGEGSLKESTKVHIINNPFLEGQSEN<br>SDWERESEGEECIAHRL |
| N44 | MERLLRKCRIRNQLVRECLAECLGVYVLILFGCGSVAQVTTSQDKKGQYLSINLG**F**ALGVTFGVFVSRGVSGAHLNPAV<br>SLSLCVLGRHPWIKLPFYVLFQVFGAFLAAATVYLQYYDAIMTYSGGQLTVTGPTATAGIFSTYPADYLSLWGGIVDQV<br>IGTAALLLCVLALGDSRNSPAPAGLEPVLVGAVVLVI**G**ISMGSNSG**Y**ALNPA**R**DLGPRLFTYIAGWGAEVFKAGGGWWW<br>VPLVAPCVGALLGTLIYELLIEVHHPPSESESQDSAQGATNQAVELEGAESDSEKPESNKQSTPMDLMERPTLGGAVCE<br>TAIEAEILGEGESVTKAHKM |

At the four pore forming sites, three residues, including ar/R residues, used for molecular weight calculation are underlined, and the ar/R residues are indicated in bold red.
