## Supplementary material for "Selectivity and evolution of Aqp10 in solute permeability influenced by pore molecular weight": Table S3

**Table S3.** Associations between the pore molecular weights in of Aqp10, Aqp10.1/1a, and Aqp10.2/2b.

| | organism | protein | position1 | position2 | position3 | position4 | mid1 | mid2 | mid3 | mid4 | MW peptide (internal) 3 aa | MW side chain sum 3 aa | MW peptide (internal) | MW side chain sum |
|---|---|---|---|---|---|---|---|---|---|---|---|---|---|---|
| | | | | | | | | | | | total 12 residues | total 12 residues | total 4 residues | total 4 residues |
| Perca flavescens AQP10.1a | Perca flavescens | AQP10.1a | AFS | ISM | GAA | ARD | F | S | A | R | 1178.36 | 506.36 | 461.53 | 237.53 |
| Perca fluviatilis AQP10.1a | Perca fluviatilis | AQP10.1a | AFS | ISM | GAA | ARD | F | S | A | R | 1178.36 | 506.36 | 461.53 | 237.53 |
| Etheostoma cragini AQP10.1a | Etheostoma cragini | AQP10.1a | AFS | ISM | GAA | ARD | F | S | A | R | 1178.36 | 506.36 | 461.53 | 237.53 |
| Etheostoma spectabile AQP10.1a | Etheostoma spectabile | AQP10.1a | AFS | ISM | GAA | ARD | F | S | A | R | 1178.36 | 506.36 | 461.53 | 237.53 |
| Sander lucioperca AQP10.1a | Sander lucioperca | AQP10.1a | AFS | ISM | GAA | ARD | F | S | A | R | 1178.36 | 506.36 | 461.53 | 237.53 |
| Cottoperca gobio AQP10.1a | Cottoperca gobio | AQP10.1a | AFS | ISM | GGA | ARD | F | S | G | R | 1164.34 | 492.34 | 447.51 | 223.51 |
| Gymnodraco acuticeps AQP10.1a | Gymnodraco acuticeps | AQP10.1a | AFS | ISM | GGA | ARD | F | S | G | R | 1164.34 | 492.34 | 447.51 | 223.51 |
| Siniperca chuatsi AQP10.1a | Siniperca chuatsi | AQP10.1a | AFS | ISM | GAA | ARD | F | S | A | R | 1178.36 | 506.36 | 461.53 | 237.53 |
| Micropterus salmoides AQP10.1a | Micropterus salmoides | AQP10.1a | AFS | ISM | GGA | ARD | F | S | G | R | 1164.34 | 492.34 | 447.51 | 223.51 |
| Micropterus dolomieu AQP10.1a | Micropterus dolomieu | AQP10.1a | AFS | ISM | GGA | ARD | F | S | G | R | 1164.34 | 492.34 | 447.51 | 223.51 |
| Dicentrarchus labrax AQP10.1a | Dicentrarchus labrax | AQP10.1a | AFS | ISM | GAA | ARD | F | S | A | R | 1178.36 | 506.36 | 461.53 | 237.53 |
| Morone saxatilis AQP10.1a | Morone saxatilis | AQP10.1a | AFS | ISM | GAA | ARD | F | S | A | R | 1178.36 | 506.36 | 461.53 | 237.53 |
| Labrus bergylta AQP10.1a | Labrus bergylta | AQP10.1a | AFS | ISM | GAA | ARD | F | S | A | R | 1178.36 | 506.36 | 461.53 | 237.53 |
| Notolabrus celidotus AQP10.1a | Notolabrus celidotus | AQP10.1a | AFS | ISM | GAA | ARD | F | S | A | R | 1178.36 | 506.36 | 461.53 | 237.53 |
| Cheilinus undulatus AQP10.1a | Cheilinus undulatus | AQP10.1a | AFS | ISM | GAA | ARD | F | S | A | R | 1178.36 | 506.36 | 461.53 | 237.53 |
| Larimichthys crocea AQP10.1a | Larimichthys crocea | AQP10.1a | AFS | ISM | GAA | ARD | F | S | A | R | 1178.36 | 506.36 | 461.53 | 237.53 |
| Acanthopagrus latus AQP10.1a | Acanthopagrus latus | AQP10.1a | AFS | ISM | GAA | ARD | F | S | A | R | 1178.36 | 506.36 | 461.53 | 237.53 |
| Sparus aurata AQP10.1a | Sparus aurata | AQP10.1a | AFS | ISM | GAA | ARD | F | S | A | R | 1178.36 | 506.36 | 461.53 | 237.53 |
| Scomber japonicus AQP10.1a | Scomber japonicus | AQP10.1a | AFS | ISM | GGA | ARD | F | S | G | R | 1164.34 | 492.34 | 447.51 | 223.51 |
| Solea senegalensis AQP10.1a | Solea senegalensis | AQP10.1a | AFS | ISM | GGA | ARD | F | S | G | R | 1164.34 | 492.34 | 447.51 | 223.51 |
| Solea solea AQP10.1a | Solea solea | AQP10.1a | AFS | ISM | GGA | ARD | F | S | G | R | 1164.34 | 492.34 | 447.51 | 223.51 |
| Scophthalmus maximus AQP10.1a | Scophthalmus maximus | AQP10.1a | AFS | ISM | GGA | ARD | F | S | G | R | 1164.34 | 492.34 | 447.51 | 223.51 |
| Paralichthys olivaceus AQP10.1a | Paralichthys olivaceus | AQP10.1a | AFS | ISM | GAA | ARD | F | S | A | R | 1178.36 | 506.36 | 461.53 | 237.53 |
| Hippoglossus stenolepis AQP10.1a | Hippoglossus stenolepis | AQP10.1a | AFS | ISM | GAA | ARD | F | S | A | R | 1178.36 | 506.36 | 461.53 | 237.53 |
| Pleuronectes platessa AQP10.1a | Pleuronectes platessa | AQP10.1a | AFS | ISV | GAA | ARD | F | S | A | R | 1146.3 | 474.3 | 461.53 | 237.53 |
| Echeneis naucrates AQP10.1a | Echeneis naucrates | AQP10.1a | AFS | ISM | GAA | ARD | F | S | A | R | 1178.36 | 506.36 | 461.53 | 237.53 |
| Seriola dumerili AQP10.1a | Seriola dumerili | AQP10.1a | AFS | ISM | GAA | ARD | F | S | A | R | 1178.36 | 506.36 | 461.53 | 237.53 |
| Seriola aureovittata AQP10.1a | Seriola aureovittata | AQP10.1a | AFS | ISM | GAA | ARD | F | S | A | R | 1178.36 | 506.36 | 461.53 | 237.53 |
| Seriola lalandi AQP10.1a | Seriola lalandi | AQP10.1a | AFS | ISM | GAA | ARD | F | S | A | R | 1178.36 | 506.36 | 461.53 | 237.53 |
| Lates calcarifer AQP10.1a | Lates calcarifer | AQP10.1a | AFS | ISM | GAA | ARD | F | S | A | R | 1178.36 | 506.36 | 461.53 | 237.53 |
| Thunnus albacares AQP10.1a | Thunnus albacares | AQP10.1a | AFS | ISM | GAA | ARD | F | S | A | R | 1178.36 | 506.36 | 461.53 | 237.53 |
| Thunnus maccoyii AQP10.1a | Thunnus maccoyii | AQP10.1a | AFS | ISM | GAA | ARD | F | S | A | R | 1178.36 | 506.36 | 461.53 | 237.53 |
| Lampris incognitus AQP10.1a | Lampris incognitus | AQP10.1a | AFS | IST | GAA | ARD | F | S | A | R | 1148.27 | 476.27 | 461.53 | 237.53 |
| Gouania willdenowi AQP10.1a | Gouania willdenowi | AQP10.1a | SFS | ISM | GAA | ARD | F | S | A | R | 1194.36 | 522.36 | 461.53 | 237.53 |
| Austrofundulus limnaeus AQP10.1a | Austrofundulus limnaeus | AQP10.1a | SFS | ISM | GAA | ARD | F | S | A | R | 1194.36 | 522.36 | 461.53 | 237.53 |
| Kryptolebias marmoratus AQP10.1a | Kryptolebias marmoratus | AQP10.1a | SFS | ISM | GAA | ARD | F | S | A | R | 1194.36 | 522.36 | 461.53 | 237.53 |
| Nematolebias whitei AQP10.1a | Nematolebias whitei | AQP10.1a | SFS | ISM | GAA | ARD | F | S | A | R | 1194.36 | 522.36 | 461.53 | 237.53 |
| Nothobranchius furzeri AQP10.1a | Nothobranchius furzeri | AQP10.1a | SFS | ISM | GAA | ARD | F | S | A | R | 1194.36 | 522.36 | 461.53 | 237.53 |
| Melanotaenia boesemani AQP10.1a | Melanotaenia boesemani | AQP10.1a | AFS | ISM | GAA | ARD | F | S | A | R | 1178.36 | 506.36 | 461.53 | 237.53 |
| Parambassis ranga AQP10.1a | Parambassis ranga | AQP10.1a | AFS | ISM | GAA | ARD | F | S | A | R | 1178.36 | 506.36 | 461.53 | 237.53 |
| Synchiropus splendidus AQP10.1a | Synchiropus splendidus | AQP10.1a | AFS | ISM | GAA | ARD | F | S | A | R | 1178.36 | 506.36 | 461.53 | 237.53 |
| Cyprinodon tularosa AQP10.1a2 | Cyprinodon tularosa | AQP10.1a2 | SFS | IGI | GAA | ARD | F | G | A | R | 1146.3 | 474.3 | 431.51 | 207.51 |
| Cyprinodon variegatus AQP10.1a2 | Cyprinodon variegatus | AQP10.1a2 | SFS | IGM | GAA | ARD | F | G | A | R | 1164.34 | 492.34 | 431.51 | 207.51 |
| Xiphophorus couchianus AQP10.1a2 | Xiphophorus couchianus | AQP10.1a2 | SFS | ISM | GAA | ARD | F | S | A | R | 1194.36 | 522.36 | 461.53 | 237.53 |
| Xiphophorus maculatus AQP10.1a2 | Xiphophorus maculatus | AQP10.1a2 | SFS | ISM | GAA | ARD | F | S | A | R | 1194.36 | 522.36 | 461.53 | 237.53 |
| Xiphophorus hellerii AQP10.1a2 | Xiphophorus hellerii | AQP10.1a2 | SFS | ISM | GAA | ARD | F | S | A | R | 1194.36 | 522.36 | 461.53 | 237.53 |
| Gambusia affinis AQP10.1a2 | Gambusia affinis | AQP10.1a2 | SFS | ISM | GAA | ARD | F | S | A | R | 1194.36 | 522.36 | 461.53 | 237.53 |
| Poeciliopsis prolifica AQP10.1a2 | Poeciliopsis prolifica | AQP10.1a2 | SFS | ISM | GAA | ARD | F | S | A | R | 1194.36 | 522.36 | 461.53 | 237.53 |
| Poecilia reticulata AQP10.1a2 | Poecilia reticulata | AQP10.1a2 | SFS | ISM | GAA | ARD | F | S | A | R | 1194.36 | 522.36 | 461.53 | 237.53 |
| Poecilia latipinna AQP10.1a2 | Poecilia latipinna | AQP10.1a2 | SFS | ISM | GAA | ARD | F | S | A | R | 1194.36 | 522.36 | 461.53 | 237.53 |
| Poecilia formosa AQP10.1a2 | Poecilia formosa | AQP10.1a2 | SFS | ISM | GAA | ARD | F | S | A | R | 1194.36 | 522.36 | 461.53 | 237.53 |
| Poecilia mexicana AQP10.1a2 | Poecilia mexicana | AQP10.1a2 | SFS | ISM | GAA | ARD | F | S | A | R | 1194.36 | 522.36 | 461.53 | 237.53 |
| Anabas testudineus AQP10.1a | Anabas testudineus | AQP10.1a | AFS | ISM | GAA | ARD | F | S | A | R | 1178.36 | 506.36 | 461.53 | 237.53 |
| Mastacembelus armatus AQP10.1a | Mastacembelus armatus | AQP10.1a | AFS | ISM | GAA | ARD | F | S | A | R | 1178.36 | 506.36 | 461.53 | 237.53 |
| Monopterus albus AQP10.1a | Monopterus albus | AQP10.1a | SFS | ISV | GGA | ARD | F | S | G | R | 1148.28 | 476.28 | 447.51 | 223.51 |
| Myripristis murdjan AQP10.1a | Myripristis murdjan | AQP10.1a | AFS | ISM | GAA | ARD | F | S | A | R | 1178.36 | 506.36 | 461.53 | 237.53 |
| Doryrhamphus excisus AQP10.1a | Doryrhamphus excisus | AQP10.1a | SFS | ISM | GAA | ARD | F | S | A | R | 1194.36 | 522.36 | 461.53 | 237.53 |
| Dunckerocampus dactyliophorus AQP10.1a | Dunckerocampus dactyliophorus | AQP10.1a | SFS | ISM | GAA | ARD | F | S | A | R | 1194.36 | 522.36 | 461.53 | 237.53 |
| Syngnathus acus AQP10.1a | Syngnathus acus | AQP10.1a | SFS | ISM | GAA | ARD | F | S | A | R | 1194.36 | 522.36 | 461.53 | 237.53 |
| Syngnathus scovelli AQP10.1a | Syngnathus scovelli | AQP10.1a | SFS | ISM | GAA | ARD | F | S | A | R | 1194.36 | 522.36 | 461.53 | 237.53 |
| Hippocampus comes AQP10.1a | Hippocampus comes | AQP10.1a | SFS | ISM | GAA | ARD | F | S | A | R | 1194.36 | 522.36 | 461.53 | 237.53 |
| Hippocampus zosterae AQP10.1a | Hippocampus zosterae | AQP10.1a | SFS | ISM | GAA | ARD | F | S | A | R | 1194.36 | 522.36 | 461.53 | 237.53 |
| Oryzias melastigma AQP10.1a | Oryzias melastigma | AQP10.1a | AFS | ISM | GGA | ARD | F | S | G | R | 1164.34 | 492.34 | 447.51 | 223.51 |
| Oryzias latipes AQP10.1a | Oryzias latipes | AQP10.1a | AFS | IAM | GGA | ARD | F | A | G | R | 1148.34 | 476.34 | 431.51 | 207.51 |
| Boleophthalmus pectinirostris AQP10.1a | Boleophthalmus pectinirostris | AQP10.1a | SFA | ISM | GAA | ARD | F | S | A | R | 1178.36 | 506.36 | 461.53 | 237.53 |
| Periophthalmus magnuspinnatus AQP10.1a | Periophthalmus magnuspinnatus | AQP10.1a | SFS | ISM | GAA | ARD | F | S | A | R | 1194.36 | 522.36 | 461.53 | 237.53 |
| Betta splendens AQP10.1a | Betta splendens | AQP10.1a | SFC | ISM | GAA | ARD | F | S | A | R | 1210.42 | 538.42 | 461.53 | 237.53 |
| Pundamilia nyererei AQP10.1a | Pundamilia nyererei | AQP10.1a | AFS | ISI | GAA | ARD | F | S | A | R | 1160.32 | 488.32 | 461.53 | 237.53 |
| Simochromis diagramma AQP10.1a | Simochromis diagramma | AQP10.1a | AFS | ISI | GAA | ARD | F | S | A | R | 1160.32 | 488.32 | 461.53 | 237.53 |
| Maylandia zebra AQP10.1a | Maylandia zebra | AQP10.1a | AFS | ISI | GAA | ARD | F | S | A | R | 1160.32 | 488.32 | 461.53 | 237.53 |
| Astatotilapia calliptera AQP10.1a | Astatotilapia calliptera | AQP10.1a | AFS | ISI | GAA | ARD | F | S | A | R | 1160.32 | 488.32 | 461.53 | 237.53 |
| Haplochromis burtoni AQP10.1a | Haplochromis burtoni | AQP10.1a | AFS | ISI | GAA | ARD | F | S | A | R | 1160.32 | 488.32 | 461.53 | 237.53 |
| Oreochromis niloticus AQP10.1a | Oreochromis niloticus | AQP10.1a | AFS | ISI | GAA | ARD | F | S | A | R | 1160.32 | 488.32 | 461.53 | 237.53 |
| Oreochromis aureus AQP10.1a | Oreochromis aureus | AQP10.1a | AFS | ISI | GAA | ARD | F | S | A | R | 1160.32 | 488.32 | 461.53 | 237.53 |
| Neolamprologus brichardi AQP10.1a | Neolamprologus brichardi | AQP10.1a | AFS | ISI | GAA | ARD | F | S | A | R | 1160.32 | 488.32 | 461.53 | 237.53 |
| Gadus chalcogrammus AQP10.1a | Gadus chalcogrammus | AQP10.1a | AFS | IAM | GGA | ARD | F | A | G | R | 1148.34 | 476.34 | 431.51 | 207.51 |
| Gadus morhua AQP10.1a | Gadus morhua | AQP10.1a | AFS | IAM | GGA | ARD | F | A | G | R | 1148.34 | 476.34 | 431.51 | 207.51 |
| Hypomesus transpacificus AQP10.1a | Hypomesus transpacificus | AQP10.1a | AFS | IAM | GGA | ARD | F | A | G | R | 1148.34 | 476.34 | 431.51 | 207.51 |
| Takifugu flavidus AQP10.1a | Takifugu flavidus | AQP10.1a | AFS | ISM | GAA | ARD | F | S | A | R | 1178.36 | 506.36 | 461.53 | 237.53 |
| Takifugu rubripes AQP10.1a | Takifugu rubripes | AQP10.1a | AFS | ISM | GAA | ARD | F | S | A | R | 1178.36 | 506.36 | 461.53 | 237.53 |
| Cyprinodon tularosa AQP10.1a1 | Cyprinodon tularosa | AQP10.1a1 | SWA | IIM | GAA | ARD | W | I | A | R | 1243.48 | 571.48 | 526.65 | 302.65 |
| Cyprinodon variegatus AQP10.1a1 | Cyprinodon variegatus | AQP10.1a1 | SWA | IIM | GAA | ARD | W | I | A | R | 1243.48 | 571.48 | 526.65 | 302.65 |
| Poeciliopsis prolifica AQP10.1a1 | Poeciliopsis prolifica | AQP10.1a1 | SWA | IVM | GAA | ARD | W | V | A | R | 1229.46 | 557.46 | 512.63 | 288.63 |
| Poecilia reticulata AQP10.1a1 | Poecilia reticulata | AQP10.1a1 | SWA | IIM | GAA | ARD | W | I | A | R | 1243.48 | 571.48 | 526.65 | 302.65 |
| Poecilia latipinna AQP10.1a1 | Poecilia latipinna | AQP10.1a1 | SWA | IIM | GAA | ARD | W | I | A | R | 1243.48 | 571.48 | 526.65 | 302.65 |
| Poecilia formosa AQP10.1a1 | Poecilia formosa | AQP10.1a1 | SWA | IIM | GAA | ARD | W | I | A | R | 1243.48 | 571.48 | 526.65 | 302.65 |
| Poecilia mexicana AQP10.1a1 | Poecilia mexicana | AQP10.1a1 | SWA | IIM | GAA | ARD | W | I | A | R | 1243.48 | 571.48 | 526.65 | 302.65 |
| Salarias fasciatus AQP10.1a | Salarias fasciatus | AQP10.1a | AFA | LSI | GCA | ARD | F | S | C | R | 1176.38 | 504.38 | 493.59 | 269.59 |
| Amphiprion ocellaris AQP10.1a | Amphiprion ocellaris | AQP10.1a | AFP | ISI | GAA | ARD | F | S | A | R | 1170.36 | 498.36 | 461.53 | 237.53 |
| Stegastes partitus AQP10.1a | Stegastes partitus | AQP10.1a | AFP | LSI | GGA | ARD | F | S | G | R | 1156.34 | 484.34 | 447.51 | 223.51 |
| Acanthochromis polyacanthus AQP10.1a | Acanthochromis polyacanthus | AQP10.1a | AFP | LSI | MGA | ARD | F | S | G | R | 1230.48 | 558.48 | 447.51 | 223.51 |
| Electrophorus electricus AQP10.2b | Electrophorus electricus | AQP10.2b | AFS | VSM | GGG | ARD | F | S | G | R | 1136.3 | 464.3 | 447.51 | 223.51 |
| Lampris incognitus AQP10.2b | Lampris incognitus | AQP10.2b | AFA | VSM | GGG | ARD | F | S | G | R | 1120.3 | 448.3 | 447.51 | 223.51 |
| Gadus chalcogrammus AQP10.1b | Gadus chalcogrammus | AQP10.1b | AFS | VSC | GGG | ARD | F | S | G | R | 1108.24 | 436.24 | 447.51 | 223.51 |
| Gadus morhua AQP10.1b | Gadus morhua | AQP10.1b | AFS | VSC | GGG | ARD | F | S | G | R | 1108.24 | 436.24 | 447.51 | 223.51 |
| Esox lucius AQP10.1a | Esox lucius | AQP10.1a | SFS | IGM | GGA | ARD | F | G | G | R | 1150.32 | 478.32 | 417.49 | 193.49 |
| Oncorhynchus mykiss AQP10.1a | Oncorhynchus mykiss | AQP10.1a | AFS | IGM | GGA | ARD | F | G | G | R | 1134.32 | 462.32 | 417.49 | 193.49 |
| Oncorhynchus kisutch AQP10.1a | Oncorhynchus kisutch | AQP10.1a | AFS | IGM | GGA | ARD | F | G | G | R | 1134.32 | 462.32 | 417.49 | 193.49 |
| Oncorhynchus tshawytscha AQP10.1a | Oncorhynchus tshawytscha | AQP10.1a | AFS | IGM | GGA | ARD | F | G | G | R | 1134.32 | 462.32 | 417.49 | 193.49 |
| Oncorhynchus keta AQP10.1a | Oncorhynchus keta | AQP10.1a | AFS | IGM | GGA | ARD | F | G | G | R | 1134.32 | 462.32 | 417.49 | 193.49 |
| Oncorhynchus nerka AQP10.1a | Oncorhynchus nerka | AQP10.1a | AFS | IGM | GGA | ARD | F | G | G | R | 1134.32 | 462.32 | 417.49 | 193.49 |
| Oncorhynchus gorbuscha AQP10.1a | Oncorhynchus gorbuscha | AQP10.1a | AFS | IGM | GGA | ARD | F | G | G | R | 1134.32 | 462.32 | 417.49 | 193.49 |
| Salvelinus alpinus AQP10.1c | Salvelinus alpinus | AQP10.1c | AFS | IGM | GYA | ARD | F | G | Y | R | 1240.44 | 568.44 | 523.61 | 299.61 |
| Salvelinus namaycush AQP10.1a | Salvelinus namaycush | AQP10.1a | AFS | IGM | GGA | ARD | F | G | G | R | 1134.32 | 462.32 | 417.49 | 193.49 |
| Salvelinus fontinalis AQP10.1a | Salvelinus fontinalis | AQP10.1a | AFS | IGM | GGA | ARD | F | G | G | R | 1134.32 | 462.32 | 417.49 | 193.49 |
| Salvelinus fontinalis AQP10.1c | Salvelinus fontinalis | AQP10.1c | AFS | IGM | GGA | ARD | F | G | G | R | 1134.32 | 462.32 | 417.49 | 193.49 |
| Brienomyrus brachyistius AQP10.1a | Brienomyrus brachyistius | AQP10.1a | GFS | ISV | GGA | ARD | F | S | G | R | 1118.26 | 446.26 | 447.51 | 223.51 |
| Paramormyrops kingsleyae AQP10.1a | Paramormyrops kingsleyae | AQP10.1a | GFS | ISV | GGA | ARD | F | S | G | R | 1118.26 | 446.26 | 447.51 | 223.51 |
| Scleropages formosus AQP10.1a | Scleropages formosus | AQP10.1a | AFC | ISL | GAA | ARD | F | S | A | R | 1176.38 | 504.38 | 461.53 | 237.53 |
| Alosa alosa AQP10.1a | Alosa alosa | AQP10.1a | AFS | ISM | GAA | ARD | F | S | A | R | 1178.36 | 506.36 | 461.53 | 237.53 |
| Alosa sapidissima AQP10.1a | Alosa sapidissima | AQP10.1a | AFS | ISM | GAA | ARD | F | S | A | R | 1178.36 | 506.36 | 461.53 | 237.53 |
| Clupea harengus AQP10.1a | Clupea harengus | AQP10.1a | AFS | IST | GGA | ARD | F | S | G | R | 1134.25 | 462.25 | 447.51 | 223.51 |
| Clupea pallasii AQP10.1a | Clupea pallasii | AQP10.1a | AFS | IST | GGA | ARD | F | S | G | R | 1134.25 | 462.25 | 447.51 | 223.51 |
| Denticeps clupeoides AQP10.1a | Denticeps clupeoides | AQP10.1a | AFS | ISI | GAA | ARD | F | S | A | R | 1160.32 | 488.32 | 461.53 | 237.53 |
| Ictalurus furcatus AQP10.1a | Ictalurus furcatus | AQP10.1a | SFA | IAT | GGA | ARD | F | A | G | R | 1118.25 | 446.25 | 431.51 | 207.51 |
| Ictalurus punctatus AQP10.1a | Ictalurus punctatus | AQP10.1a | SFA | IAI | GGA | ARD | F | A | G | R | 1130.3 | 458.3 | 431.51 | 207.51 |
| Pangasianodon hypophthalmus AQP10.1a | Pangasianodon hypophthalmus | AQP10.1a | SFA | IGM | GGA | ARD | F | G | G | R | 1134.32 | 462.32 | 417.49 | 193.49 |
| Silurus meridionalis AQP10.1a | Silurus meridionalis | AQP10.1a | SFA | IAM | GGA | ARD | F | A | G | R | 1148.34 | 476.34 | 431.51 | 207.51 |
| Tachysurus fulvidraco AQP10.1a | Tachysurus fulvidraco | AQP10.1a | SFA | IAM | GGA | ARD | F | A | G | R | 1148.34 | 476.34 | 431.51 | 207.51 |
| Hemibagrus wyckioides AQP10.1a | Hemibagrus wyckioides | AQP10.1a | SFA | IAM | GGA | ARD | F | A | G | R | 1148.34 | 476.34 | 431.51 | 207.51 |
| Astyanax mexicanus AQP10.1a | Astyanax mexicanus | AQP10.1a | SFA | IAM | GGA | ARD | F | A | G | R | 1148.34 | 476.34 | 431.51 | 207.51 |
| Pygocentrus nattereri AQP10.1a | Pygocentrus nattereri | AQP10.1a | SFA | IAM | GGA | ARD | F | A | G | R | 1148.34 | 476.34 | 431.51 | 207.51 |
| Colossoma macropomum AQP10.1a | Colossoma macropomum | AQP10.1a | SFA | IAM | GGA | ARD | F | A | G | R | 1148.34 | 476.34 | 431.51 | 207.51 |
| Electrophorus electricus AQP10.1a | Electrophorus electricus | AQP10.1a | SFA | IAM | GGA | ARD | F | A | G | R | 1148.34 | 476.34 | 431.51 | 207.51 |
| Chanos chanos AQP10.1a | Chanos chanos | AQP10.1a | AFS | ISV | GAA | ARD | F | S | A | R | 1146.3 | 474.3 | 461.53 | 237.53 |
| Puntigrus tetrazona AQP10.1a | Puntigrus tetrazona | AQP10.1a | AFS | ISM | GAA | ARD | F | S | A | R | 1178.36 | 506.36 | 461.53 | 237.53 |
| Sinocyclocheilus anshuiensis AQP10.1a | Sinocyclocheilus anshuiensis | AQP10.1a | AFS | ISM | GAA | ARD | F | S | A | R | 1178.36 | 506.36 | 461.53 | 237.53 |
| Sinocyclocheilus rhinocerous AQP10.1a | Sinocyclocheilus rhinocerous | AQP10.1a | AFS | ISM | GAA | ARD | F | S | A | R | 1178.36 | 506.36 | 461.53 | 237.53 |
| Sinocyclocheilus grahami AQP10.1a | Sinocyclocheilus grahami | AQP10.1a | AFS | ISM | GAA | ARD | F | S | A | R | 1178.36 | 506.36 | 461.53 | 237.53 |
| Onychostoma macrolepis AQP10.1a | Onychostoma macrolepis | AQP10.1a | AFS | ISM | GAA | ARD | F | S | A | R | 1178.36 | 506.36 | 461.53 | 237.53 |
| Carassius gibelio AQP10.1a2 | Carassius gibelio | AQP10.1a2 | AFS | ISM | GAA | ARD | F | S | A | R | 1178.36 | 506.36 | 461.53 | 237.53 |
| Carassius gibelio AQP10.1a1 | Carassius gibelio | AQP10.1a1 | AFS | ISM | GAA | ARD | F | S | A | R | 1178.36 | 506.36 | 461.53 | 237.53 |
| Megalobrama amblycephala AQP10.1a | Megalobrama amblycephala | AQP10.1a | AFS | ISM | GAA | ARD | F | S | A | R | 1178.36 | 506.36 | 461.53 | 237.53 |
| Danio aesculapii AQP10.1a | Danio aesculapii | AQP10.1a | AFS | ISI | GAA | ARD | F | S | A | R | 1160.32 | 488.32 | 461.53 | 237.53 |
| Danio rerio AQP10.1a | Danio rerio | AQP10.1a | AFS | ISI | GAA | ARD | F | S | A | R | 1160.32 | 488.32 | 461.53 | 237.53 |
| Pimephales promelas AQP10.1a | Pimephales promelas | AQP10.1a | AFS | ISM | GAA | ARD | F | S | A | R | 1178.36 | 506.36 | 461.53 | 237.53 |
| Rhinichthys klamathensis AQP10.1a | Rhinichthys klamathensis | AQP10.1a | AFS | ISM | GAA | ARD | F | S | A | R | 1178.36 | 506.36 | 461.53 | 237.53 |
| Megalops cyprinoides AQP10.1a | Megalops cyprinoides | AQP10.1a | AFS | ISM | GAA | ARD | F | S | A | R | 1178.36 | 506.36 | 461.53 | 237.53 |
| Lepisosteus oculatus AQP10.1 | Lepisosteus oculatus | AQP10.1 | SFA | ISM | GAA | ARD | F | S | A | R | 1178.36 | 506.36 | 461.53 | 237.53 |
| Erpetoichthys calabaricus AQP10.1 | Erpetoichthys calabaricus | AQP10.1 | AFA | ISM | GAA | ARD | F | S | A | R | 1162.36 | 490.36 | 461.53 | 237.53 |
| Polypterus senegalus AQP10.1 | Polypterus senegalus | AQP10.1 | AFA | ISM | GAA | ARD | F | S | A | R | 1162.36 | 490.36 | 461.53 | 237.53 |

| | | | | | | | | | | | | | | |
|---|---|---|---|---|---|---|---|---|---|---|---|---|---|---|
| Acipenser ruthenus AQP10.1a | Acipenser ruthenus | AQP10.1a | AFA | IGM | GGA | ARD | F | G | G | R | 1118.32 | 446.32 | 417.49 | 193.49 |
| Polyodon spathula AQP10.1b | Polyodon spathula | AQP10.1b | AFA | IGM | GGA | ARD | F | G | G | R | 1118.32 | 446.32 | 417.49 | 193.49 |
| Acipenser ruthenus AQP10.2b | Acipenser ruthenus | AQP10.2b | AFA | IGL | GYA | ARD | F | G | Y | R | 1206.4 | 534.4 | 523.61 | 299.61 |
| Acipenser ruthenus AQP10.2a | Acipenser ruthenus | AQP10.2a | ASA | IGL | GYA | ARD | S | G | Y | R | 1146.3 | 474.3 | 463.51 | 239.51 |
| Polyodon spathula AQP10.2b | Polyodon spathula | AQP10.2b | AFA | IGL | GYA | ARD | F | G | Y | R | 1206.4 | 534.4 | 523.61 | 299.61 |
| Erpetoichthys calabaricus AQP10.2 | Erpetoichthys calabaricus | AQP10.2 | AFA | IGI | GYA | ARD | F | G | Y | R | 1206.4 | 534.4 | 523.61 | 299.61 |
| Polypterus senegalus AQP10.2 | Polypterus senegalus | AQP10.2 | AFA | IGI | GYA | ARD | F | G | Y | R | 1206.4 | 534.4 | 523.61 | 299.61 |
| Alosa alosa AQP10.2a | Alosa alosa | AQP10.2a | AFS | IGL | GYP | ARD | F | G | Y | R | 1248.44 | 576.44 | 523.61 | 299.61 |
| Alosa sapidissima AQP10.2a | Alosa sapidissima | AQP10.2a | AFS | IGL | GYP | ARD | F | G | Y | R | 1248.44 | 576.44 | 523.61 | 299.61 |
| Clupea harengus AQP10.2a | Clupea harengus | AQP10.2a | AFA | IGL | GYP | ARD | F | G | Y | R | 1232.44 | 560.44 | 523.61 | 299.61 |
| Clupea pallasii AQP10.2a | Clupea pallasii | AQP10.2a | AFA | IGL | GYP | ARD | F | G | Y | R | 1232.44 | 560.44 | 523.61 | 299.61 |
| Lepisosteus oculatus AQP10.2 | Lepisosteus oculatus | AQP10.2 | AFA | IGI | GYP | ARD | F | G | Y | R | 1232.44 | 560.44 | 523.61 | 299.61 |
| Anguilla anguilla AQP10.2b2 | Anguilla anguilla | AQP10.2b2 | GFA | IGM | GGA | ARD | F | G | G | R | 1104.3 | 432.3 | 417.49 | 193.49 |
| Anguilla anguilla AQP10.2b1 | Anguilla anguilla | AQP10.2b1 | GFA | IGM | GGA | ARD | F | G | G | R | 1104.3 | 432.3 | 417.49 | 193.49 |
| Anguilla anguilla AQP10.2b3 | Anguilla anguilla | AQP10.2b3 | GFA | IGI | GYA | ARD | F | G | Y | R | 1192.38 | 520.38 | 523.61 | 299.61 |
| Megalops cyprinoides AQP10.2b | Megalops cyprinoides | AQP10.2b | GFA | IGI | GYA | ARD | F | G | Y | R | 1192.38 | 520.38 | 523.61 | 299.61 |
| Carassius gibelio AQP10.2b | Carassius gibelio | AQP10.2b | GFA | IGI | GYA | ARD | F | G | Y | R | 1192.38 | 520.38 | 523.61 | 299.61 |
| Sinocyclocheilus rhinocerous AQP10.2b2 | Sinocyclocheilus rhinocerous | AQP10.2b2 | GFA | IGI | GYA | ARD | F | G | Y | R | 1192.38 | 520.38 | 523.61 | 299.61 |
| Onychostoma macrolepis AQP10.2b | Onychostoma macrolepis | AQP10.2b | GFA | IGI | GYA | ARD | F | G | Y | R | 1192.38 | 520.38 | 523.61 | 299.61 |
| Sinocyclocheilus anshuiensis AQP10.2b | Sinocyclocheilus anshuiensis | AQP10.2b | GFA | IGI | GYA | ARD | F | G | Y | R | 1192.38 | 520.38 | 523.61 | 299.61 |
| Sinocyclocheilus rhinocerous AQP10.2b1 | Sinocyclocheilus rhinocerous | AQP10.2b1 | GFA | IGI | GYA | ARD | F | G | Y | R | 1192.38 | 520.38 | 523.61 | 299.61 |
| Sinocyclocheilus grahami AQP10.2b1 | Sinocyclocheilus grahami | AQP10.2b1 | GFA | IGI | GYA | ARD | F | G | Y | R | 1192.38 | 520.38 | 523.61 | 299.61 |
| Danio aesculapii AQP10.2b | Danio aesculapii | AQP10.2b | GFA | IGI | GYA | ARD | F | G | Y | R | 1192.38 | 520.38 | 523.61 | 299.61 |
| Danio rerio AQP10.2b | Danio rerio | AQP10.2b | GFA | IGI | GYA | ARD | F | G | Y | R | 1192.38 | 520.38 | 523.61 | 299.61 |
| Megalobrama amblycephala AQP10.2b | Megalobrama amblycephala | AQP10.2b | GFA | IGI | GYA | ARD | F | G | Y | R | 1192.38 | 520.38 | 523.61 | 299.61 |
| Rhinichthys klamathensis AQP10.2b | Rhinichthys klamathensis | AQP10.2b | GFA | IGI | GYA | ARD | F | G | Y | R | 1192.38 | 520.38 | 523.61 | 299.61 |
| Pygocentrus nattereri AQP10.2b | Pygocentrus nattereri | AQP10.2b | GFA | IGI | GYA | ARD | F | G | Y | R | 1192.38 | 520.38 | 523.61 | 299.61 |
| Colossoma macropomum AQP10.2b | Colossoma macropomum | AQP10.2b | GFA | IGI | GYA | ARD | F | G | Y | R | 1192.38 | 520.38 | 523.61 | 299.61 |
| Astyanax mexicanus AQP10.2b | Astyanax mexicanus | AQP10.2b | GFA | IGI | GYA | ARD | F | G | Y | R | 1192.38 | 520.38 | 523.61 | 299.61 |
| Pangasianodon hypophthalmus AQP10.2b | Pangasianodon hypophthalmus | AQP10.2b | GFA | IGV | GYA | ARD | F | G | Y | R | 1178.36 | 506.36 | 523.61 | 299.61 |
| Ictalurus furcatus AQP10.2b | Ictalurus furcatus | AQP10.2b | GFA | IGV | GYA | ARD | F | G | Y | R | 1178.36 | 506.36 | 523.61 | 299.61 |
| Ictalurus punctatus AQP10.2b | Ictalurus punctatus | AQP10.2b | GFA | IGV | GYA | ARD | F | G | Y | R | 1178.36 | 506.36 | 523.61 | 299.61 |
| Silurus meridionalis AQP10.2b | Silurus meridionalis | AQP10.2b | GFA | IGV | GYA | ARD | F | G | Y | R | 1178.36 | 506.36 | 523.61 | 299.61 |
| Tachysurus fulvidraco AQP10.2b | Tachysurus fulvidraco | AQP10.2b | AFA | IGL | GYA | ARD | F | G | Y | R | 1206.4 | 534.4 | 523.61 | 299.61 |
| Hemibagrus wyckioides AQP10.2b | Hemibagrus wyckioides | AQP10.2b | GFA | IGV | GYA | ARD | F | G | Y | R | 1178.36 | 506.36 | 523.61 | 299.61 |
| Alosa alosa AQP10.2b | Alosa alosa | AQP10.2b | GFA | IGV | GYA | ARD | F | G | Y | R | 1178.36 | 506.36 | 523.61 | 299.61 |
| Alosa sapidissima AQP10.2b | Alosa sapidissima | AQP10.2b | GFA | IGV | GYA | ARD | F | G | Y | R | 1178.36 | 506.36 | 523.61 | 299.61 |
| Clupea harengus AQP10.2b | Clupea harengus | AQP10.2b | GFA | IGV | GYA | ARD | F | G | Y | R | 1178.36 | 506.36 | 523.61 | 299.61 |
| Clupea pallasii AQP10.2b | Clupea pallasii | AQP10.2b | GFA | IGV | GYA | ARD | F | G | Y | R | 1178.36 | 506.36 | 523.61 | 299.61 |
| Hypomesus transpacificus AQP10.2b | Hypomesus transpacificus | AQP10.2b | GFA | IGV | GYS | ARD | F | G | Y | R | 1194.36 | 522.36 | 523.61 | 299.61 |
| Esox lucius AQP10.2b | Esox lucius | AQP10.2b | GFA | IGI | GYS | ARD | F | G | Y | R | 1208.38 | 536.38 | 523.61 | 299.61 |
| Salvelinus fontinalis AQP10.2b | Salvelinus fontinalis | AQP10.2b | GFA | IGV | GYA | ARD | F | G | Y | R | 1178.36 | 506.36 | 523.61 | 299.61 |
| Salvelinus namaycush AQP10.2b | Salvelinus namaycush | AQP10.2b | GFA | IGV | GYA | ARD | F | G | Y | R | 1178.36 | 506.36 | 523.61 | 299.61 |
| Salvelinus alpinus AQP10.2b | Salvelinus alpinus | AQP10.2b | GFA | IGV | GGA | ARD | F | G | G | R | 1072.24 | 400.24 | 417.49 | 193.49 |
| Oncorhynchus mykiss AQP10.2b | Oncorhynchus mykiss | AQP10.2b | GFA | IGV | GYA | ARD | F | G | Y | R | 1178.36 | 506.36 | 523.61 | 299.61 |
| Oncorhynchus tshawytscha AQP10.2b | Oncorhynchus tshawytscha | AQP10.2b | GFA | IGV | GYA | ARD | F | G | Y | R | 1178.36 | 506.36 | 523.61 | 299.61 |
| Oncorhynchus gorbuscha AQP10.2b | Oncorhynchus gorbuscha | AQP10.2b | GFA | IGV | GYA | ARD | F | G | Y | R | 1178.36 | 506.36 | 523.61 | 299.61 |
| Oncorhynchus nerka AQP10.2b | Oncorhynchus nerka | AQP10.2b | GFA | IGV | GYA | ARD | F | G | Y | R | 1178.36 | 506.36 | 523.61 | 299.61 |
| Oncorhynchus kisutch AQP10.2b | Oncorhynchus kisutch | AQP10.2b | GFA | IGV | GYA | ARD | F | G | Y | R | 1178.36 | 506.36 | 523.61 | 299.61 |
| Oncorhynchus keta AQP10.2b | Oncorhynchus keta | AQP10.2b | GFA | IGV | GYA | ARD | F | G | Y | R | 1178.36 | 506.36 | 523.61 | 299.61 |
| Gadus chalcogrammus AQP10.2b | Gadus chalcogrammus | AQP10.2b | GFA | IGV | GYA | ARD | F | G | Y | R | 1178.36 | 506.36 | 523.61 | 299.61 |
| Gadus morhua AQP10.2b | Gadus morhua | AQP10.2b | GFA | IGV | GYA | ARD | F | G | Y | R | 1178.36 | 506.36 | 523.61 | 299.61 |
| Myripristis murdjan AQP10.2b | Myripristis murdjan | AQP10.2b | GFA | IGI | GYA | ARD | F | G | Y | R | 1192.38 | 520.38 | 523.61 | 299.61 |
| Lampris incognitus AQP10.2b2 | Lampris incognitus | AQP10.2b2 | GFA | IGV | GYA | ARD | F | G | Y | R | 1178.36 | 506.36 | 523.61 | 299.61 |
| Thalassophryne amazonica AQP10.2b | Thalassophryne amazonica | AQP10.2b | GFA | IGI | GYA | ARD | F | G | Y | R | 1192.38 | 520.38 | 523.61 | 299.61 |
| Boleophthalmus pectinirostris AQP10.2b | Boleophthalmus pectinirostris | AQP10.2b | GFA | IGV | GYA | ARD | F | G | Y | R | 1178.36 | 506.36 | 523.61 | 299.61 |
| Periophthalmus magnuspinnatus AQP10.2b | Periophthalmus magnuspinnatus | AQP10.2b | GFA | IGV | GYA | ARD | F | G | Y | R | 1178.36 | 506.36 | 523.61 | 299.61 |
| Synchiropus splendidus AQP10.2b | Synchiropus splendidus | AQP10.2b | GFA | IGA | GYA | ARD | F | G | Y | R | 1150.3 | 478.3 | 523.61 | 299.61 |
| Cheilinus undulatus AQP10.2b | Cheilinus undulatus | AQP10.2b | GFA | IGI | GYA | ARD | F | G | Y | R | 1192.38 | 520.38 | 523.61 | 299.61 |
| Takifugu flavidus AQP10.2b | Takifugu flavidus | AQP10.2b | GFS | IGI | GYA | ARD | F | G | Y | R | 1208.38 | 536.38 | 523.61 | 299.61 |
| Takifugu rubripes AQP10.2b | Takifugu rubripes | AQP10.2b | GFS | IGI | GYA | ARD | F | G | Y | R | 1208.38 | 536.38 | 523.61 | 299.61 |
| Gouania willdenowi AQP10.2b | Gouania willdenowi | AQP10.2b | GFA | IGI | GYA | ARD | F | G | Y | R | 1192.38 | 520.38 | 523.61 | 299.61 |
| Salarias fasciatus AQP10.2b | Salarias fasciatus | AQP10.2b | GFA | IGI | GYA | ARD | F | G | Y | R | 1192.38 | 520.38 | 523.61 | 299.61 |
| Perca flavescens AQP10.2b | Perca flavescens | AQP10.2b | GFA | IGI | GYA | ARD | F | G | Y | R | 1192.38 | 520.38 | 523.61 | 299.61 |
| Perca fluviatilis AQP10.2b | Perca fluviatilis | AQP10.2b | GFA | IGI | GYA | ARD | F | G | Y | R | 1192.38 | 520.38 | 523.61 | 299.61 |
| Sander lucioperca AQP10.2b | Sander lucioperca | AQP10.2b | GFA | IGI | GYA | ARD | F | G | Y | R | 1192.38 | 520.38 | 523.61 | 299.61 |
| Etheostoma cragini AQP10.2b | Etheostoma cragini | AQP10.2b | GFA | IGI | GYA | ARD | F | G | Y | R | 1192.38 | 520.38 | 523.61 | 299.61 |
| Etheostoma spectabile AQP10.2b | Etheostoma spectabile | AQP10.2b | GFA | IGI | GYA | ARD | F | G | Y | R | 1192.38 | 520.38 | 523.61 | 299.61 |
| Dicentrarchus labrax AQP10.2b | Dicentrarchus labrax | AQP10.2b | GFA | IGI | GYA | ARD | F | G | Y | R | 1192.38 | 520.38 | 523.61 | 299.61 |
| Morone saxatilis AQP10.2b | Morone saxatilis | AQP10.2b | GFA | IGI | GYA | ARD | F | G | Y | R | 1192.38 | 520.38 | 523.61 | 299.61 |
| Siniperca chuatsi AQP10.2b | Siniperca chuatsi | AQP10.2b | GFA | IGI | GYA | ARD | F | G | Y | R | 1192.38 | 520.38 | 523.61 | 299.61 |
| Micropterus salmoides AQP10.2b | Micropterus salmoides | AQP10.2b | GFA | IGI | GYA | ARD | F | G | Y | R | 1192.38 | 520.38 | 523.61 | 299.61 |
| Micropterus dolomieu AQP10.2b | Micropterus dolomieu | AQP10.2b | GFA | IGI | GYA | ARD | F | G | Y | R | 1192.38 | 520.38 | 523.61 | 299.61 |
| Thunnus albacares AQP10.2b | Thunnus albacares | AQP10.2b | GFA | IGI | GYA | ARD | F | G | Y | R | 1192.38 | 520.38 | 523.61 | 299.61 |
| Thunnus maccoyii AQP10.2b | Thunnus maccoyii | AQP10.2b | GFA | IGI | GYA | ARD | F | G | Y | R | 1192.38 | 520.38 | 523.61 | 299.61 |
| Scomber japonicus AQP10.2b | Scomber japonicus | AQP10.2b | GFA | IGV | GYA | ARD | F | G | Y | R | 1178.36 | 506.36 | 523.61 | 299.61 |
| Lates calcarifer AQP10.2b | Lates calcarifer | AQP10.2b | GFA | IGI | GYA | ARD | F | G | Y | R | 1192.38 | 520.38 | 523.61 | 299.61 |
| Echeneis naucrates AQP10.2b | Echeneis naucrates | AQP10.2b | GFA | IGV | GYA | ARD | F | G | Y | R | 1178.36 | 506.36 | 523.61 | 299.61 |
| Seriola dumerili AQP10.2b | Seriola dumerili | AQP10.2b | GFA | IGI | GYA | ARD | F | G | Y | R | 1192.38 | 520.38 | 523.61 | 299.61 |
| Seriola aureovittata AQP10.2b | Seriola aureovittata | AQP10.2b | GFA | IGI | GYA | ARD | F | G | Y | R | 1192.38 | 520.38 | 523.61 | 299.61 |
| Seriola lalandi AQP10.2b | Seriola lalandi | AQP10.2b | GFA | IGI | GYA | ARD | F | G | Y | R | 1192.38 | 520.38 | 523.61 | 299.61 |
| Gymnodraco acuticeps AQP10.2b | Gymnodraco acuticeps | AQP10.2b | GFA | IGI | GYA | ARD | F | G | Y | R | 1192.38 | 520.38 | 523.61 | 299.61 |
| Betta splendens AQP10.2b | Betta splendens | AQP10.2b | GFA | IGI | GYA | ARD | F | G | Y | R | 1192.38 | 520.38 | 523.61 | 299.61 |
| Scophthalmus maximus AQP10.2b | Scophthalmus maximus | AQP10.2b | GFA | IGI | GYA | ARD | F | G | Y | R | 1192.38 | 520.38 | 523.61 | 299.61 |
| Monopterus albus AQP10.2b | Monopterus albus | AQP10.2b | GFA | IGV | GYP | AGD | F | G | Y | G | 1105.27 | 433.27 | 424.48 | 200.48 |
| Anabas testudineus AQP10.2b | Anabas testudineus | AQP10.2b | GFA | IGI | GYA | ARD | F | G | Y | R | 1192.38 | 520.38 | 523.61 | 299.61 |
| Cottoperca gobio AQP10.2b | Cottoperca gobio | AQP10.2b | GFA | IGI | GYA | ARD | F | G | Y | R | 1192.38 | 520.38 | 523.61 | 299.61 |
| Mastacembelus armatus AQP10.2b | Mastacembelus armatus | AQP10.2b | GFA | IGI | GYA | ARD | F | G | Y | R | 1192.38 | 520.38 | 523.61 | 299.61 |
| Channa argus AQP10.2b | Channa argus | AQP10.2b | GFA | IGI | GYA | ARD | F | G | Y | R | 1192.38 | 520.38 | 523.61 | 299.61 |
| Paralichthys olivaceus AQP10.2b | Paralichthys olivaceus | AQP10.2b | GFA | IGI | GYA | ARD | F | G | Y | R | 1192.38 | 520.38 | 523.61 | 299.61 |
| Hippoglossus stenolepis AQP10.2b | Hippoglossus stenolepis | AQP10.2b | GFA | IGI | GYA | ARD | F | G | Y | R | 1192.38 | 520.38 | 523.61 | 299.61 |
| Pleuronectes platessa AQP10.2b | Pleuronectes platessa | AQP10.2b | GFA | IGI | GYA | ARD | F | G | Y | R | 1192.38 | 520.38 | 523.61 | 299.61 |
| Solea senegalensis AQP10.2b | Solea senegalensis | AQP10.2b | GFA | IGI | GYA | ARD | F | G | Y | R | 1192.38 | 520.38 | 523.61 | 299.61 |
| Solea solea AQP10.2b | Solea solea | AQP10.2b | GFA | IGI | GYA | ARD | F | G | Y | R | 1192.38 | 520.38 | 523.61 | 299.61 |
| Parambassis ranga AQP10.2b | Parambassis ranga | AQP10.2b | GFA | IGV | GYA | ARD | F | G | Y | R | 1178.36 | 506.36 | 523.61 | 299.61 |
| Labrus bergylta AQP10.2b | Labrus bergylta | AQP10.2b | GFA | IGI | GYA | ARD | F | G | Y | R | 1192.38 | 520.38 | 523.61 | 299.61 |
| Notolabrus celidotus AQP10.2b | Notolabrus celidotus | AQP10.2b | GFA | IGI | GYA | ARD | F | G | Y | R | 1192.38 | 520.38 | 523.61 | 299.61 |
| Astatotilapia calliptera AQP10.2b | Astatotilapia calliptera | AQP10.2b | GFA | IGI | GYA | ARD | F | G | Y | R | 1192.38 | 520.38 | 523.61 | 299.61 |
| Maylandia zebra AQP10.2b | Maylandia zebra | AQP10.2b | GFA | IGI | GYA | ARD | F | G | Y | R | 1192.38 | 520.38 | 523.61 | 299.61 |
| Pundamilia nyererei AQP10.2b | Pundamilia nyererei | AQP10.2b | GFA | IGI | GYA | ARD | F | G | Y | R | 1192.38 | 520.38 | 523.61 | 299.61 |
| Haplochromis burtoni AQP10.2b | Haplochromis burtoni | AQP10.2b | GFA | IGI | GYA | ARD | F | G | Y | R | 1192.38 | 520.38 | 523.61 | 299.61 |
| Simochromis diagramma AQP10.2b | Simochromis diagramma | AQP10.2b | GFA | IGI | GYA | ARD | F | G | Y | R | 1192.38 | 520.38 | 523.61 | 299.61 |
| Neolamprologus brichardi AQP10.2b | Neolamprologus brichardi | AQP10.2b | GFA | IGI | GYA | ARD | F | G | Y | R | 1192.38 | 520.38 | 523.61 | 299.61 |
| Oreochromis niloticus AQP10.2b | Oreochromis niloticus | AQP10.2b | GFA | IGI | GYA | ARD | F | G | Y | R | 1192.38 | 520.38 | 523.61 | 299.61 |
| Oreochromis aureus AQP10.2b | Oreochromis aureus | AQP10.2b | GFA | IGI | GYA | ARD | F | G | Y | R | 1192.38 | 520.38 | 523.61 | 299.61 |
| Larimichthys crocea AQP10.2b | Larimichthys crocea | AQP10.2b | GFA | IGV | GYA | ARD | F | G | Y | R | 1178.36 | 506.36 | 523.61 | 299.61 |
| Acanthopagrus latus AQP10.2b | Acanthopagrus latus | AQP10.2b | GFA | IGV | GYA | ARD | F | G | Y | R | 1178.36 | 506.36 | 523.61 | 299.61 |
| Sparus aurata AQP10.2b | Sparus aurata | AQP10.2b | GFA | IGV | GYA | ARD | F | G | Y | R | 1178.36 | 506.36 | 523.61 | 299.61 |
| Amphiprion ocellaris AQP10.2b | Amphiprion ocellaris | AQP10.2b | GFA | LGI | GYA | ARD | F | G | Y | R | 1192.38 | 520.38 | 523.61 | 299.61 |
| Acanthochromis polyacanthus AQP10.2b | Acanthochromis polyacanthus | AQP10.2b | GFA | LGI | GYA | ARD | F | G | Y | R | 1192.38 | 520.38 | 523.61 | 299.61 |
| Stegastes partitus AQP10.2b | Stegastes partitus | AQP10.2b | GFA | IGV | GYA | ARD | F | G | Y | R | 1178.36 | 506.36 | 523.61 | 299.61 |
| Oryzias melastigma AQP10.2b | Oryzias melastigma | AQP10.2b | GFA | IGV | GYA | ARD | F | G | Y | R | 1178.36 | 506.36 | 523.61 | 299.61 |
| Oryzias latipes AQP10.2b | Oryzias latipes | AQP10.2b | GFA | IGV | GYA | ARD | F | G | Y | R | 1178.36 | 506.36 | 523.61 | 299.61 |
| Melanotaenia boesemani AQP10.2b | Melanotaenia boesemani | AQP10.2b | GFA | IGV | GYA | ARD | F | G | Y | R | 1178.36 | 506.36 | 523.61 | 299.61 |
| Nematolebias whitei AQP10.2b | Nematolebias whitei | AQP10.2b | SFA | IGI | GYP | ARD | F | G | Y | R | 1248.44 | 576.44 | 523.61 | 299.61 |
| Austrofundulus limnaeus AQP10.2b | Austrofundulus limnaeus | AQP10.2b | GFA | IGI | GYP | ARD | F | G | Y | R | 1218.42 | 546.42 | 523.61 | 299.61 |
| Kryptolebias marmoratus AQP10.2b | Kryptolebias marmoratus | AQP10.2b | GFA | IGI | GYA | ARD | F | G | Y | R | 1192.38 | 520.38 | 523.61 | 299.61 |
| Nothobranchius furzeri AQP10.2b | Nothobranchius furzeri | AQP10.2b | GFA | IGV | GYA | ARD | F | G | Y | R | 1178.36 | 506.36 | 523.61 | 299.61 |
| Cyprinodon tularosa AQP10.2b | Cyprinodon tularosa | AQP10.2b | GFA | IGM | GYA | ARD | F | G | Y | R | 1210.42 | 538.42 | 523.61 | 299.61 |
| Cyprinodon variegatus AQP10.2b | Cyprinodon variegatus | AQP10.2b | GFA | IGM | GYA | ARD | F | G | Y | R | 1210.42 | 538.42 | 523.61 | 299.61 |
| Poeciliopsis prolifica AQP10.2b | Poeciliopsis prolifica | AQP10.2b | GFA | IGV | GYA | ARD | F | G | Y | R | 1178.36 | 506.36 | 523.61 | 299.61 |
| Poecilia formosa AQP10.2b | Poecilia formosa | AQP10.2b | GFA | IGV | GYA | ARD | F | G | Y | R | 1178.36 | 506.36 | 523.61 | 299.61 |
| Poecilia latipinna AQP10.2b | Poecilia latipinna | AQP10.2b | GFA | IGV | GYA | ARD | F | G | Y | R | 1178.36 | 506.36 | 523.61 | 299.61 |
| Poecilia mexicana AQP10.2b | Poecilia mexicana | AQP10.2b | GFA | IGV | GYA | ARD | F | G | Y | R | 1178.36 | 506.36 | 523.61 | 299.61 |
| Poecilia reticulata AQP10.2b | Poecilia reticulata | AQP10.2b | GFA | IGV | GYA | ARD | F | G | Y | R | 1178.36 | 506.36 | 523.61 | 299.61 |
| Gambusia affinis AQP10.2b | Gambusia affinis | AQP10.2b | GFA | IGV | GYA | ARD | F | G | Y | R | 1178.36 | 506.36 | 523.61 | 299.61 |
| Xiphophorus hellerii AQP10.2b | Xiphophorus hellerii | AQP10.2b | GFA | IGV | GYA | ARD | F | G | Y | R | 1178.36 | 506.36 | 523.61 | 299.61 |
| Xiphophorus couchianus AQP10.2b | Xiphophorus couchianus | AQP10.2b | GFA | IGV | GYA | ARD | F | G | Y | R | 1178.36 | 506.36 | 523.61 | 299.61 |
| Xiphophorus maculatus AQP10.2b | Xiphophorus maculatus | AQP10.2b | GFA | IGV | GYA | ARD | F | G | Y | R | 1178.36 | 506.36 | 523.61 | 299.61 |
| Latimeria chalumnae AQP10 | Latimeria chalumnae | AQP10 | ACG | IGM | GYA | ARD | C | G | Y | R | 1166.38 | 494.38 | 479.57 | 255.57 |
| Protopterus annectens AQP10 | Protopterus annectens | AQP10 | GAA | LGT | GYA | TRD | A | G | Y | R | 1134.26 | 462.26 | 447.51 | 223.51 |
| Rhinatrema bivittatum AQP10 | Rhinatrema bivittatum | AQP10 | AGA | IGL | GYP | ARD | G | G | Y | R | 1142.32 | 470.32 | 433.49 | 209.49 |
| Xenopus laevis AQP10 | Xenopus laevis | AQP10 | AGA | IGL | SYP | TRD | G | G | Y | R | 1202.37 | 530.37 | 433.49 | 209.49 |
| Xenopus tropicalis AQP10 | Xenopus tropicalis | AQP10 | AGA | IGL | GYP | TRD | G | G | Y | R | 1172.35 | 500.35 | 433.49 | 209.49 |
| Bufo bufo AQP10 | Bufo bufo | AQP10 | AGA | IGL | GFP | SRD | G | G | F | R | 1142.32 | 470.32 | 417.49 | 193.49 |
| Gallus gallus AQP10 | Gallus gallus | AQP10 | AGA | IEV | GCP | ARD | G | E | C | R | 1140.32 | 468.32 | 445.51 | 221.51 |
| Falco cherrug AQP10 | Falco cherrug | AQP10 | AGA | IEV | GCP | ARD | G | E | C | R | 1140.32 | 468.32 | 445.51 | 221.51 |
| Crocodylus porosus AQP10 | Crocodylus porosus | AQP10 | AGA | LGL | SCA | ARD | G | G | C | R | 1086.26 | 414.26 | 373.45 | 149.45 |
| Gekko japonicus AQP10 | Gekko japonicus | AQP10 | ASG | LSL | SCA | ARD | S | S | C | R | 1132.28 | 460.28 | 433.49 | 209.49 |
| Sarcophilus harrisii AQP10 | Sarcophilus harrisii | AQP10 | AGA | IGL | GYP | ARD | G | G | Y | R | 1142.32 | 470.32 | 433.49 | 209.49 |
| Monodelphis domestica AQP10 | Monodelphis domestica | AQP10 | AGS | IGL | GYP | ARD | G | G | Y | R | 1158.32 | 486.32 | 433.49 | 209.49 |
| Oryctolagus cuniculus AQP10 | Oryctolagus cuniculus | AQP10 | GGA | IGL | GYP | ARD | G | G | Y | R | 1128.3 | 456.3 | 433.49 | 209.49 |
| Sus scrofa AQP10 | Sus scrofa | AQP10 | AGA | IGL | GYP | ARD | G | G | Y | R | 1142.32 | 470.32 | 433.49 | 209.49 |
| Tupaia chinensis AQP10 | Tupaia chinensis | AQP10 | AGS | IGL | GFP | ARD | G | G | F | R | 1142.32 | 470.32 | 417.49 | 193.49 |
| Dasypus novemcinctus AQP10 | Dasypus novemcinctus | AQP10 | AAS | IGL | GFP | ARD | A | G | F | R | 1156.34 | 484.34 | 431.51 | 207.51 |

| | | | | | | | | | | | | | |
|---|---|---|---|---|---|---|---|---|---|---|---|---|---|
| Choloepus didactylus AQP10 | Choloepus didactylus | AQP10 | AGS | IGL | GFP | ARD | G | G | F | R | 1142.32 | 470.32 | 417.49 | 193.49 |
| Carlito syrichta AQP10 | Carlito syrichta | AQP10 | AGS | IGL | GFP | ARD | G | G | F | R | 1142.32 | 470.32 | 417.49 | 193.49 |
| Cavia porcellus AQP10 | Cavia porcellus | AQP10 | AGS | IGL | GFP | ARD | G | G | F | R | 1142.32 | 470.32 | 417.49 | 193.49 |
| Homo sapiens AQP10 | homo sapiens | AQP10 | AGS | LGL | GIP | ARD | G | G | I | R | 1108.3 | 436.3 | 383.47 | 159.47 |
| Loxodonta africana AQP10 | Loxodonta africana | AQP10 | AGS | LAL | GLP | ARD | G | A | L | R | 1122.32 | 450.32 | 397.49 | 173.49 |
| Canis lupus familiaris AQP10 | Canis lupus familiaris | AQP10 | AAS | IGL | GFP | ARD | A | G | F | R | 1156.34 | 484.34 | 431.51 | 207.51 |
| Felis catus AQP10 | Felis catus | AQP10 | AGS | IGL | GFP | ARD | G | G | F | R | 1142.32 | 470.32 | 417.49 | 193.49 |
