## Supplementary material for "Selectivity and evolution of Aqp10 in solute permeability influenced by pore molecular weight": Table S4

**Table S4.** List of primers used for the infusion cloning and site-directed mutagenesis of Aqp10s.

| Species | Gene | Accession | Remarks | Direction | Sequence (5' to 3') |
|---|---|---|---|---|---|
| | *N22* | | In-Fusion cloning | Fw | gcagatcaattccccATGGAGAAACTC |
| | | | | Rv | agaattcggatccccTCAAAGCCGATG |
| | | | Site-directed mutagenesis (A205Y) | Fw | GAGCAGCAATTGTGGATACGCTATTAACCCTGC |
| | | | | Rv | GCAGGGTTAATAGCGTATCCACAATTGCTGCTC |
| | *N42* | | In-Fusion cloning | Fw | gcagatcaattccccATGGAGAAAGTGCGTCGC |
| | | | | Rv | agaattcggatccccTCATAACCGATGAGCAATGC |
| | | | Site-directed mutagenesis (F56G) | Fw | GTCCATCAATCTCGCAGGTGCTCTTGGAGTCAC |
| | | | | Rv | GTGACTCCAAGAGCACCTGCGAGATTGATGGAC |
| | | | Site-directed mutagenesis (Y205A) | Fw | GGCTCCAATTGCGGGGCTGCAATCAATCCTGC |
| | | | | Rv | GCAGGATTGATTGCAGCCCCGCAATTGGAGCC |
| | *N44* | | In-Fusion cloning | Fw | gcagatcaattccccATGGAACGGTTACTCCGC |
| | | | | Rv | TCACATTTTGTGGGCTTTAGTCACACTC (1st),<br>agaattcggatccccTCACATTTTGTGGGCTTTAG (2nd) |
| Gray bichir | *aqp10.1* | XM_039751835.1 | Site-directed mutagenesis (S196G) | Fw | GTGCTAGGAATAGGTATGTCGATGAGCTCAAACTG |
| | | | | Rv | CAGTTTGAGCTCATCGACATACCTATTCCTAGCAC |
| | | | Site-directed mutagenesis (A205Y) | Fw | GAGCTCAAACTGTGGTTATGCCATCAATCCAGCCC |
| | | | | Rv | GGGCTGGATTGATGGCATAACCACAGTTTGAGCTC |
| Gray bichir | *aqp10.2* | XM_039751813.1 | Site-directed mutagenesis (F56G) | Fw | CTTTCCATCAACCTGGCAGGTGCTATTGGCATCAC |
| | | | | Rv | GTGATGCCAATAGCACCTGCCAGGTTGATGGAAAG |
| | | | Site-directed mutagenesis (Y205A) | Fw | GGGATCCAATTGTGGTGCTGCTATCAATCCAGCCC |
| | | | | Rv | GGGCTGGATTGATAGCAGCACCACAATTGGATCCC |
| Zebrafish | *aqp10.1a* | NM_001002349.1 | Site-directed mutagenesis (S192G) | Fw | GTTCTTGGGATTGGCATCTCAATGTCGGCTAATTG |
| | | | | Rv | CAATTAGCCGACATTGAGATGCCAATCCCAAGAAC |
| | | | Site-directed mutagenesis (A205Y) | Fw | GTCGGCTAATTGTGGATATGCCATAAATCCAGCAC |
| | | | | Rv | GTGCTGGATTTATGGCATATCCACAATTAGCCGAC |
| Zebrafish | *aqp10.2b* | XM_005159392.4 | Site-directed mutagenesis (F56G) | Fw | GTCAATCAACCTGGGCGGCGCACTGGGAACCAC |
| | | | | Rv | GTGGTTCCCAGTGCGCCGCCCAGGTTGATTGAC |
| | | | In-Fusion cloning | Fw | gcagatcaattccccATGGACCGTCTGCTGAGG |
| | | | *DreAqp10.2 Y205A were chemically synthesized and subcloned. | Rv | agaattcggatccccTTACCCTACGTCTGCTGAGG |
| Human | *aqp10* | NP_536354 | Site-directed mutagenesis (G62F) | Fw | CACCATGTTTCTGGCTTTCTCTCTGGCCGTTACGA |
| | | | | Rv | TCGTAACGGCCAGAGAGAAAGCCAGAAACATGGTG |
| African clawed frog | *aqp10.L* | XM_041573749.1 | Site-directed mutagenesis (G86F) | Fw | CTGGCATTCGCTATTGCGGTGGTTCTG |
| | | | | Rv | AATAGCGAATGCCAGGTACATGCAGAAG |
