## Supplementary material for "Selectivity and evolution of Aqp10 in solute permeability influenced by pore molecular weight": Fig. S1

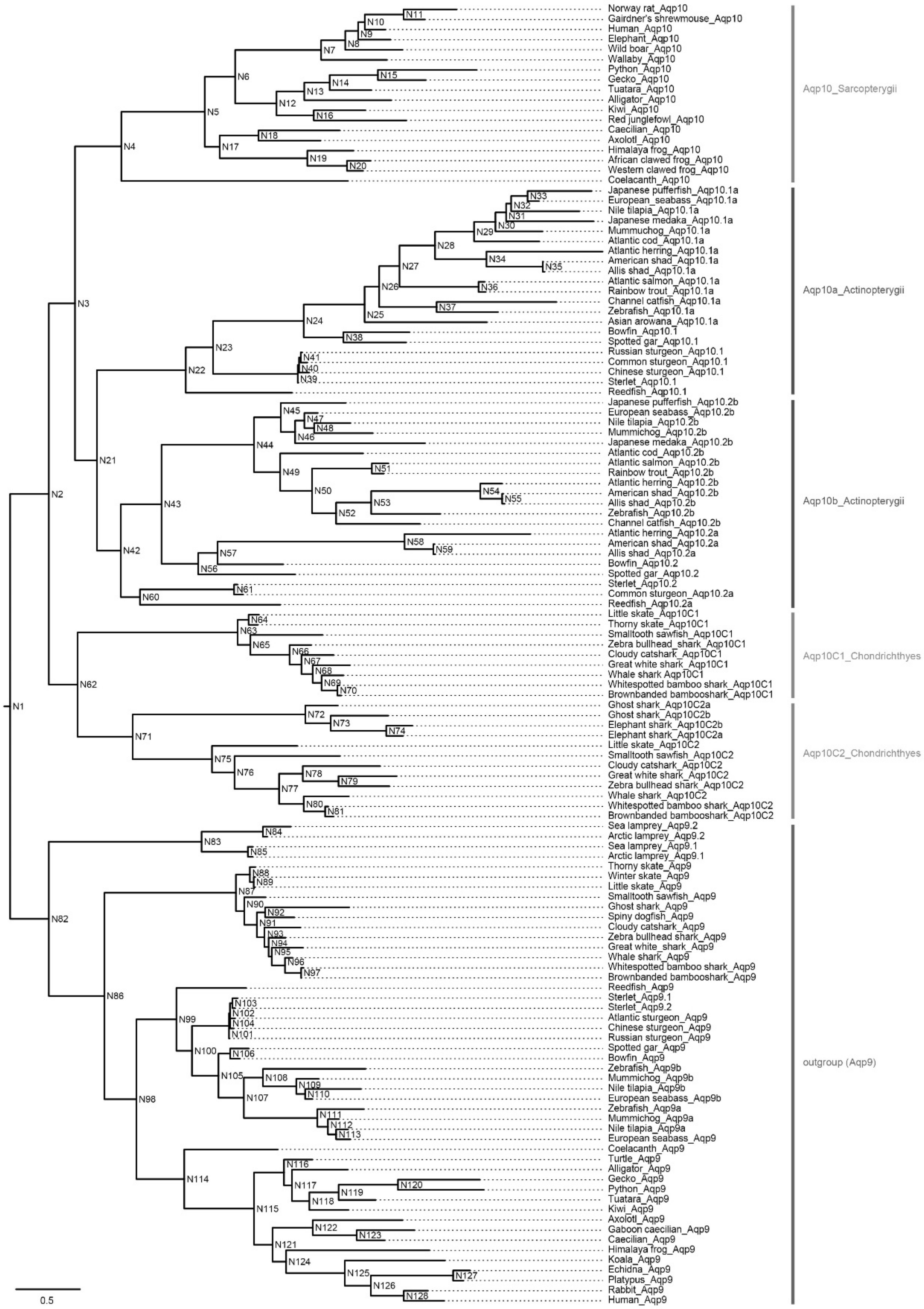


**Fig. S1.** Ancestral Aqp10 sequences. Ancestral sequences of Aqp10s estimated via phylogenetic analysis using FastML. Each node in the phylogenetic tree has a number beginning with N.
