## Supplementary material for "Selectivity and evolution of Aqp10 in solute permeability influenced by pore molecular weight": Fig. S2

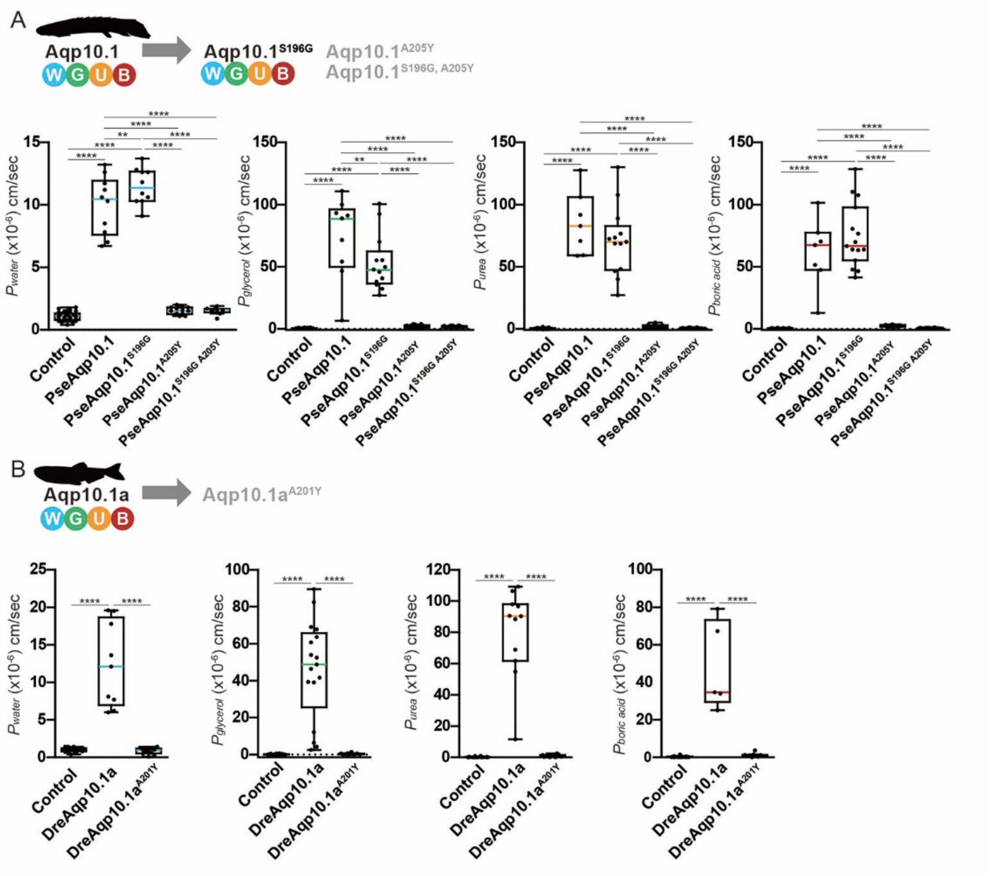

**Fig. S2.** Water and solute permeabilities of gray bichir and zebrafish Aqp10.1/1a and their mutants. Water ($P_{water}$), glycerol ($P_{glycerol}$), urea ($P_{urea}$), and boric acid ($P_{boric\ acid}$) permeabilities of gray bichir Aqp10.1 (PseAqp10.1) (*A*) and zebrafish Aqp10.1 (DreAqp10.1a) (*B*). Change in the volume of oocytes expressing each Aqp10 was compared with that of control oocytes. Values are represented as interquartile ranges from the 25$^{th}$ to 75$^{th}$ percentiles (box), range (whiskers), outliers (>1.5 × interquartile range above the upper quartile), and median (line in the box). Mean values, standard deviations, and total numbers of assayed oocytes are summarized in Table 1. Statistical significance was assessed via a one-way analysis of variance (ANOVA) followed by Tukey's test (****$p < 0.0001$; ***$p < 0.001$; **$p < 0.01$). The sample number is listed in Table S1.
