## Supplementary material for "Selectivity and evolution of Aqp10 in solute permeability influenced by pore molecular weight": Fig. S3

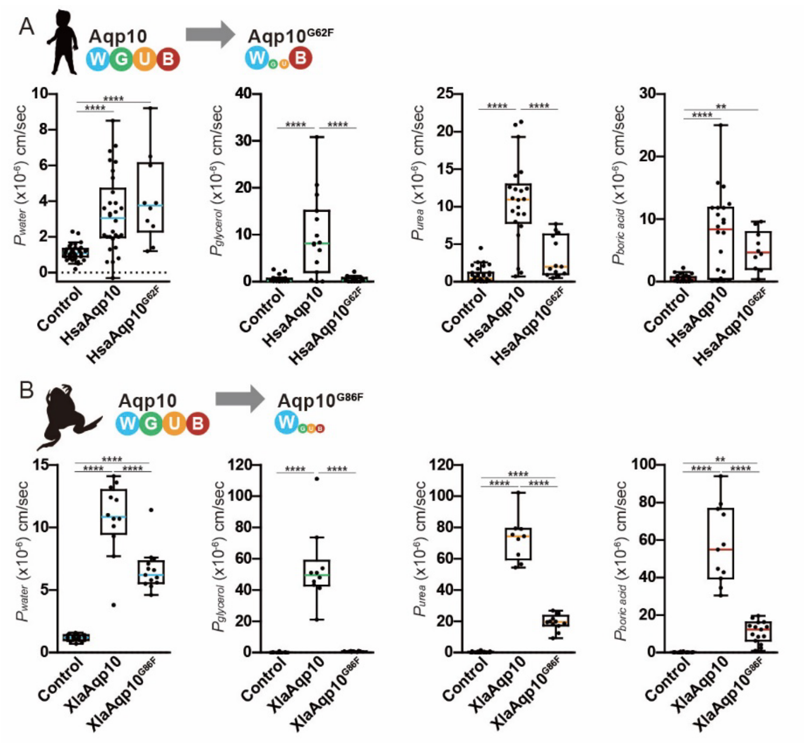


**Fig. S3.** Water and solute permeabilities of human and African clawed frog Aqp10s and their mutants. Water ($P_{\mathrm{water}}$), glycerol ($P_{\mathrm{glycerol}}$), urea ($P_{\mathrm{urea}}$), and boric acid ($P_{\mathrm{boric\ acid}}$) permeabilities of human Aqp10 (HsaAqp10) (A) and African clawed frog Aqp10 (XlaAqp10) (B). Data were plotted using box-and-whisker plots (median: 25–75 percentile; 1.5 × interquartile). Statistical significance was assessed via a one-way analysis of variance (ANOVA) followed by Tukey's test (****$p$ < 0.0001; **$p$ < 0.01). The sample number is listed in Table S1.
